# A neurovascular template guides the spatial and functional compartmentalization of the adrenal gland

**DOI:** 10.1101/2025.07.22.666120

**Authors:** Alessia Motta, Santo Diprima, Aurora Badaloni, Ganesh Parameshwar Bhat, Elena Ioannou, Chiara Malpighi, Ilaria Brambilla, Chiara Saulle, Alessandro Sessa, Christiana Ruhrberg, Dario Bonanomi

## Abstract

The vasculature adapts to tissue demands, but whether it can instruct spatial tissue organization remains unclear. In the developing adrenal gland, we uncover a neurovascular mechanism that actively establishes and preserves compartment boundaries between cortex and medulla. Peripheral nerves secrete Semaphorin3C, signaling through PlexinD1 on endothelial cells to locally antagonize VEGF-driven angiogenesis from cortical cells, sculpting distinct vascular domains that guide hormone-producing cells to their correct territories. Disruption of this balance —via denervation or loss of Semaphorin3C-PlexinD1 signaling— leads to ectopic vascularization of the medulla, which adopts a cortex-like vascular network. This vascular remodeling enables cortical cells invasion of medullary territories, blurring compartment boundaries and triggering a phagocytic macrophage response that reflects pathological hijacking of a postnatal morphogenesis program. Our findings reveal that region-specific vascular scaffolds, shaped by neurovascular cues, serve as instructive templates for organ architecture. Failure of neurovascular signaling can thus trigger a cascade of structural collapse that undermines tissue integrity and homeostasis, driving pathological remodeling.

## Introduction

With few exceptions, every organ and tissue in the body features pervasive networks of nerves and blood vessels that are essential for organ physiology and homeostasis. Innervation and vascularization typically proceed in parallel during organogenesis, as nerve fibers and vessels grow into developing tissues, engaging in reciprocal crosstalk that often results in anatomical matching between the two systems. Beyond their mutual interactions, organotypic neurovascular patterning is guided by parenchymal cells to ensure these networks meet tissue-specific demands (Miao *et al*, 2025). However, it remains less clear to what extent nerves and vessels simply follow the specific developmental blueprint of each organ or whether they also contribute to shaping it. This raises the question of whether neurovascular components actively instruct organogenesis, beyond merely responding to parenchymal cues.

New roles for peripheral innervation in organ assembly and homeostasis, beyond their canonical functions in neuronal circuit wiring and neurotransmission, are increasingly recognized. Nerve-derived signals have been shown to modulate immune responses (Ueno *et al*, 2016; Hanoun *et al*, 2015), regulate stem cell pool maintenance in adult tissues (Hanoun *et al*, 2015), and shape vascular networks during development, regeneration, and disease (Gutnick *et al*, 2011; Li *et al*, 2013; James & Mukouyama, 2011; Carmeliet & Tessier-Lavigne, 2005; Vieira *et al*, 2022; Martins *et al*, 2022; Cattin *et al*, 2015). Further underscoring a role in tissue morphogenesis, axons can serve as migratory routes for progenitor cells (Furlan *et al*, 2017; Kastriti *et al*, 2019; Erickson *et al*, 2024), but they can also facilitate cancer cell spread during metastasis (Hanoun *et al*, 2015; Wang *et al*, 2020). Likewise, the acquisition of organ-specific structural and functional features by vascular beds is no longer seen merely as a byproduct of their circulatory role in delivering oxygen and nutrients. Instead, the emerging molecular heterogeneity of endothelial and perivascular cells underlies the specification of vascular niches that release angiocrine factors and interact with the surrounding microenvironment to regulate tissue differentiation and morphogenesis across multiple organs systems, including bone, liver, lung, and kidney (Bishop *et al*, 2023; Mizrachi *et al*, 1990; Stucker *et al*, 2021; Chen *et al*, 2021). Although increasing evidence suggests that nerves and vessels interact with developing tissues in complex ways, how their coordinated integration influence organ architecture remains elusive.

Neuroendocrine organs such as the adrenal gland offer a compelling context to study neurovascular integration, given their reliance on tightly coordinated neural signaling and hormone release into the bloodstream. Functioning as a regulatory hub, the adrenal gland integrates sympathetic and pituitary inputs to maintain homeostasis under both basal and stress conditions. Its physiological output depends on the interplay between two anatomically and functionally distinct compartments: the outer cortex, which regulates metabolism, immune function, and blood pressure by releasing steroid hormones in response to circulating factors like pituitary Adrenocorticotropic Hormone (ACTH); and the inner neuroendocrine medulla, where chromaffin cells secrete catecholamines upon direct stimulation from visceral motor neurons in the spinal cord, triggering the “fight-or-flight” response —the near-instantaneous sequence of whole-body changes enabling rapid survival reactions to life-threatening situations. This compartmentalized organization is gradually acquired during embryonic development as visceral motor axons that reach the mesoderm-derived adrenocortical primordium (Luo *et al*, 1994) guide and deliver nerve-associated Schwann cell precursors (SCPs) of neural crest origin, which differentiate into medullary chromaffin cells (Furlan *et al*, 2017). While initially intermingled, cortical and medullary cells progressively segregate into distinct zones (Wood *et al*, 2013). The proportionate development and communication between adrenal cortex and medulla are critical to prevent hormonal imbalance, which would significantly affect the development of multiple organs, parturition, early postnatal life and adult stress response (Hillman *et al*, 2012; Pignatti *et al*, 2023; Nurse *et al*, 2018; Ream *et al*, 2008; Thomas *et al*, 1995; Zhou *et al*, 1995; Fitch *et al*, 2012; Huang *et al*, 2012; Sato *et al*, 2023). Deregulated growth and disrupted hormone output commonly occur in tumors arising from either compartment, including adrenocortical carcinomas and adenomas, or pheochromocytomas and neuroblastomas that originate from chromaffin cell lineages in the medulla (Bechmann *et al*, 2021; Kastriti *et al*, 2020; Huber *et al*, 2018; Maris *et al*, 2007; Matthay *et al*, 2016). The coordinated assembly and structural integrity of the adrenal gland depend on reciprocal interactions between cortical and chromaffin cells, or their progenitors, during development (Bechmann *et al*, 2021; Kastriti *et al*, 2020); yet, other components of the adrenal microenvironment likely contribute as well, though their roles remain to be defined.

As one of the most highly vascularized organs in the body, the adrenal gland relies on a precise spatial correspondence between vascular architecture and the organization of hormone-producing cells in the cortex and medulla to sustain its dual endocrine functions (Kikuta & Murakami, 1982; Stucker *et al*, 2021). Endothelial cells with specialized structural and molecular characteristics give rise to vascular networks tailored to the unique functional needs of each region. In the cortex, a dense network of fenestrated capillaries closely associates with steroidogenic cells, enabling efficient uptake of systemic inductive cues and timely hormone secretion. By contrast, the medulla is supplied by sparser sinusoidal vessels that surround innervated clusters of chromaffin cells (Kikuta & Murakami, 1984). Although the mechanisms establishing this regional vascular heterogeneity remain poorly understood, studies have shown that ACTH, a key driver of fetal adrenal cortex growth, induces the expression the pro-angiogenic signals VEGF (Vascular Endothelial Growth Factor and ANG-2 (Angiopoietin-2) in cortical cells, promoting vascular expansion in parallel with organ development (Vittet *et al*, 2000; Thomas *et al*, 2004; Ishimoto *et al*, 2006; Shifren *et al*, 1998). Elevated VEGF expression in the cortex likely underlies the higher vascular density and permeability observed in this region compared to the medulla (Thomas *et al*, 2003; Bumbasirevic *et al*, 1990). While the stereotyped distribution of blood vessels and nerves across endocrine compartments clearly supports adrenal function, it also suggests that region-specific neurovascular interactions with parenchymal cells may actively guide adrenal patterning during organogenesis.

We show that regional vascular patterning, shaped by nerve-derived factors, is a key determinant of adrenal compartmentalization. Sema3C released from motor axons and associated SCPs signals through endothelial Plexin-D1 to organize blood vessels in the medulla. When this pathway is disrupted, a cortex-like vascular network forms in the medulla. VEGF-expressing cortical cells, which display strong vessel affinity, track these ectopic routes into medullary territories, further amplifying mispatterned angiogenesis and promoting aberrant cortical expansion. This maladaptive feedback loop progressively erodes the cortex–medulla boundary, linking neurovascular dysregulation to structural disorganization of the adrenal gland.

## Results

### Spatiotemporal coordination of neurovascular patterning and adrenal gland compartmentalization

To investigate the dynamic interactions among cellular populations during adrenal gland organogenesis, we established a developmental timeline of cortex–medulla compartmentalization in relation to vascular ingrowth and innervation in mouse embryos. At E11.5, the adrenal anlagen, located ventrolateral to the dorsal aorta, consisted of adrenocortical progenitors of mesodermal origin that expressed the nuclear receptor Steroidogenic Factor 1 (SF1) —a key regulator of steroidogenic cell identity, survival, and activity (Luo *et al*, 1994; Crawford *et al*, 1997; Doghman *et al*, 2007) (Figure 1A). At this early stage, the adrenal primordium was already vascularized, with CD31-labeled blood vessels extending into the SF1^+^ cell mass (Figure 1A, G, G’). Shortly thereafter, between E12.0 and E12.5, the adrenal medulla began to form as visceral motor axons —visualized using the *MN^(218-2)^::GFP* reporter, active in spinal motor neurons (Amin *et al*, 2015)— invaded the adrenal primordium delivering SOX10^+^ SCPs (Figures 1H-I’, O). These SCPs acquired chromaffin cell identity, marked by the expression of the early sympathoadrenal specification factor PHOX2B and terminal differentiation enzyme tyrosine hydroxylase (TH). At early stages (E12.5–E13.5), cortical (SF1⁺) and medullary (TH⁺) cells were intermingled (Figure 1B, C), but progressively resolved into distinct compartments, which were clearly delineated by late gestation (E15.5–E18.5) (Figure 1D, F).

**Figure 1:**
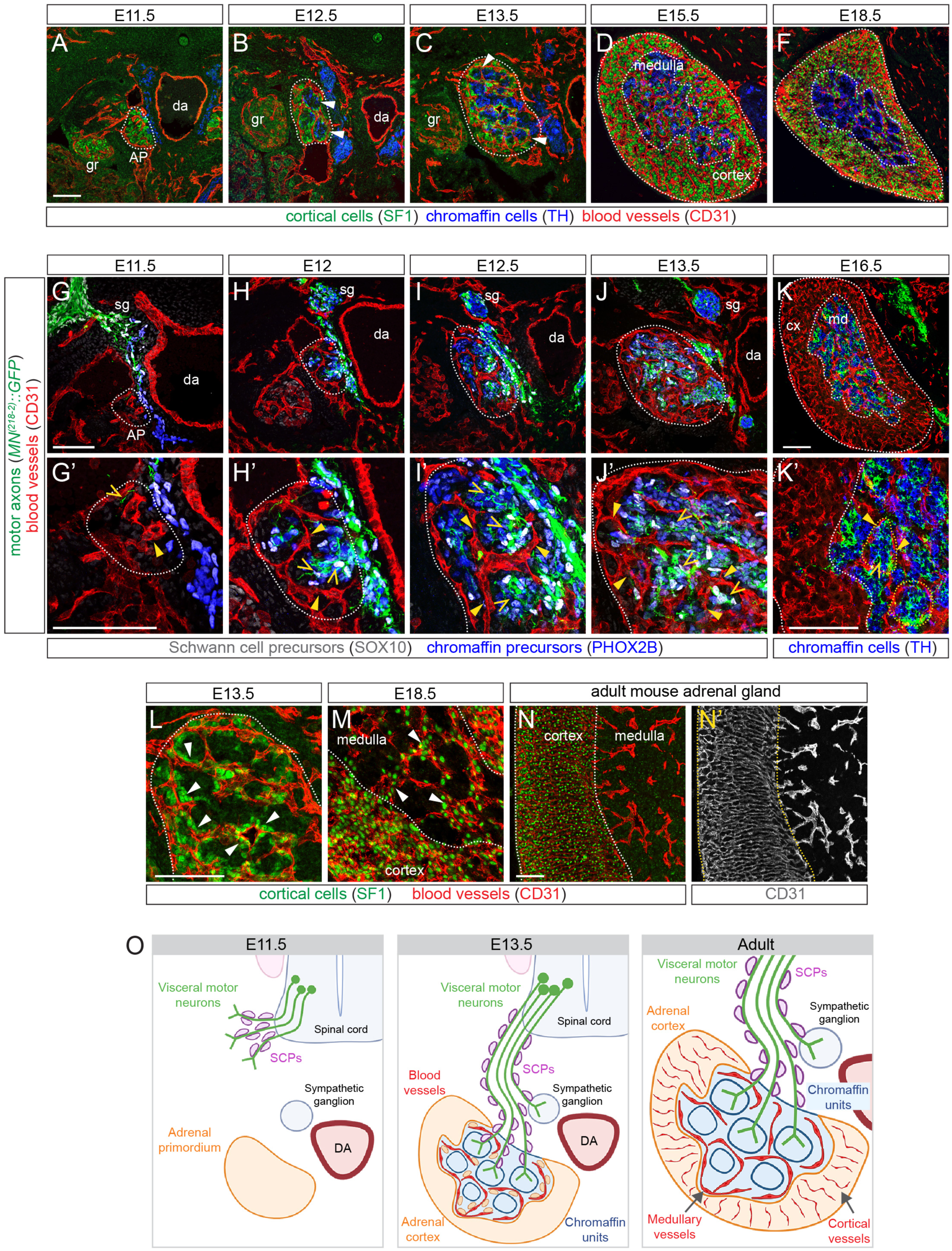
Time-course of adrenal gland development. **(A –F)** Immunostaining for cortical marker SF1 (green), TH for medullary chromaffin cells (blue) and CD31 for blood vessels (red). **(G – K’)** Immunostaining for endothelial CD31 (red), SCP marker SOX10 (grey) and the chromaffin precursor marker PHOX2B (E11.5 – E13.5) or differentiated chromaffin cell marker TH (E16.5) in blue. Motor axon terminals are labelled by *MN^(218-2)^::GFP* (green). G’ – K’ shows magnified views from G – K. **(L – N’)** Association of cortical cells (SF1^+^) with vessels at E13.5 (L), E18.5 (M), and in adult adrenal gland. L and M are magnified from C and F, respectively. CD31 staining is shown separately in N’. **(O)** Schematic of AG development showing nerve-assisted SCP invasion of the adrenal primordium, progressive cortex-medulla segregation and compartment-dependent vascular maturation. Scale bars: 100 µm. AP: adrenal primordium; cx: cortex; da: dorsal aorta; gr: genital ridge; md: medulla; SCPs: Schwann cell precursors; sg: sympathetic ganglion.

This compartmentalization was accompanied by a pronounced reconfiguration of both blood vessels and motor axons in the developing adrenal gland. The rudimentary vascular network of the adrenal primordium at E11.5 progressively increased in topological complexity over time. However, its spatial pattern remained relatively homogeneous until E13.5, when the segregation between cortical and medullary cell populations had not yet begun (Figure 1G-J’ and Figure S1A-D). After E15.5, regional differences in vascular organization became apparent: the cortex exhibited a denser, more irregular vascular meshwork, while the medulla developed a sparser, honeycomb-like pattern of larger vessels that outlined chromaffin cell clusters (Figure 1K, K’ and Figure S1E). This regional specialization anticipated the distinct architectural and functional features of the adult adrenal vasculature, where tightly packed, radially arranged fenestrated vessels in the cortex contrast with the sparse, sinusoidal vessels of the medulla (Kikuta & Murakami, 1982, 1984) (Figure 1N, N’ and Figure S1F-G’). Similarly, motor axons entering the adrenal anlagen were initially broadly distributed but gradually became spatially aligned with chromaffin cell clusters encircled by blood vessels, thereby forming discrete functional units by E16.5 (Figure 1G-K’, O and Figure S1A’-E’). This precise anatomical congruency between innervation and vasculature likely facilitates efficient neurogenic regulation of chromaffin cell activity and hormone release into the circulation.

Notably, SF1⁺ cortical cells were consistently associated with blood vessels from the early stages of adrenal development (Figure 1A-D). Consequently, the vessels coursing through the developing medulla were ensheathed by cortical cells, suggesting that they may have a higher affinity for blood vessels than chromaffin cells (Figure 1L). As cortex–medulla segregation progressed, blood vessels in the medulla remained partially covered by cortical cells surrounding the emerging chromaffin clusters (Figure 1M). It was only postnatally that cortical cells fully retracted from the medulla, resulting in the sharply defined cortical and medullary boundary characteristic of the adult gland (Wood *et al*, 2013; Kinga *et al*, 2009; Chang *et al*, 2011) (Figure 1N, N, O). The notion of a differential affinity for vessels was further supported by co-culture assays in which cortical and medullary cells from dissociated E14.5 adrenal glands were combined with fetal human endothelial cells (HUVECs). Initially intermixed, these cell types self-organized over time: cortical cells were in contact with endothelial cells, eventually forming an intervening layer between medullary clusters and the endothelium (Figure S1H), reminiscent of the *in vivo* tissue configuration (Figure S1H’).

This developmental analysis reveals that neurovascular patterning is tightly coupled to the establishment of adrenal tissue architecture, raising the question of whether these processes are functionally interdependent and collectively required for proper organ morphogenesis.

### Early visceral motor innervation shapes adrenal vascular patterning

Given the close coordination of neurovascular remodeling and adrenal compartmentalization, we sought to determine whether visceral motor innervation contributes to the establishment of adrenal vascular architecture. To investigate this, we evaluated the effects of denervation on adrenal angiogenesis by examining *Olig2^Cre^; R26-DTA^LSL^* embryos, in which spinal motor neurons and their projections are ablated via expression of Diphtheria toxin A (DTA) driven by *Olig2^Cre^* in motor neuron progenitors (Figure 2A). By E11.5, these embryos displayed a virtually complete loss of motor innervation, as evidenced by the absence of *MN^(218-2)^::GFP*-labeled fibers (Figure S2A). Since visceral motor nerves serve as migratory tracks for SCPs, denervation resulted in a severe depletion of the medullary chromaffin cells, consistent with previous studies (Furlan *et al*, 2017; Lumb *et al*, 2018) (Figure 2B). However, the ∼65% reduction in medulla area in *Olig2^Cre^; R26-DTA^LSL^* embryos was not accompanied by a proportionate decrease in adrenal gland size, which was reduced by only 10–15% in mutants suggesting that the cortex, or possibly the surrounding capsule, may have undergone a corresponding expansion (Figure 2C). In motor neuron-ablated embryos, we observed a striking remodeling of the adrenal vascular network starting at E12.5 and continuing through later stages. Blood vessels grew into the region normally occupied by the medulla, forming a denser and more compact meshwork (Figure 2D, D’, G, G’). Unlike in control embryos, where larger vascular loops typically circumscribed chromaffin cell clusters, the mutant vasculature formed smaller, tightly packed vascular compartments —most lacking medullary cells— with reduced intervascular spacing (Figure 2E-L). These ectopic vessels expressed normal levels of VEGFR2/KDR and TIE2, associated with angiogenesis and vascular stabilization, respectively, and were covered by pericytes, consistent with a typical maturation program despite their aberrant location (Figure S2B-E). Notably, by E18.5, the adrenal vasculature had lost its regional cortical-medullary distinction, adopting a cortex-like pattern that extended across the entire gland, suggesting that vascular architecture adapts to the absence of medullary tissue (Figure 2M).

**Figure 2:**
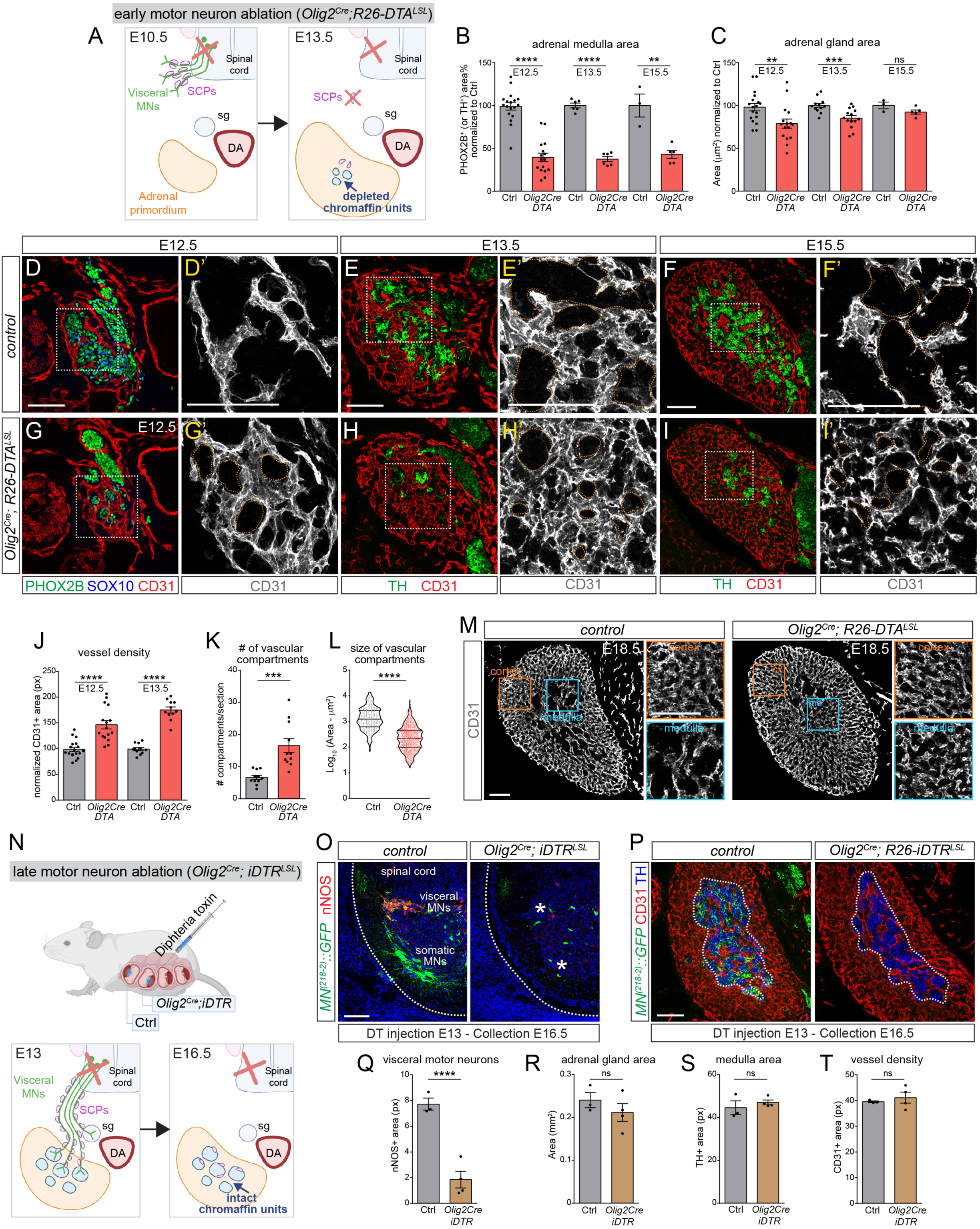
Early motor innervation controls vascularization of the adrenal gland. **(A)** Schematic of genetic motor neuron (MN) ablation with Diphtheria toxin A (DTA) expression in MN progenitors using the *Olig2^Cre^*. **(B – C)** Size of the adrenal medulla (B), identified as either PHOX2B^+^ or TH^+^ cell area, and total adrenal gland area (C) in *Olig2^Cre^; R26-DTA^LSL^* and controls at E12.5, E13.5 and E15.5. **(D – I’)** Adrenal gland sections from *Olig2^Cre^; R26-DTA^LSL^* embryos (G – I’) and controls (D – F’) at E12.5 (D, D’, G, G’), E13.5 (E, E’, H, H’) and E15.5 (F, F’, I, I’). CD31 identifies vessels (red), TH and PHOX2B label chromaffin cells (green), SCPs are labelled with SOX10 (blue). Magnified views of CD31 (grey) from the boxed areas are shown in the lower panels (D’, E’, F’, G’, H’, I’). Adrenal compartments encircled by adrenal vessels are outlined (orange). **(J)** Adrenal vessel density in *Olig2^Cre^; R26-DTA^LSL^* and control embryos at E12.5 and E13.5. **(K)** Number of adrenal-vascular compartments at E13.5. **(L)** Size distribution of the size of adrenal-vascular compartments at E13.5. Dots correspond to individual compartments. **(M)** Adrenal gland sections of E18.5 *Olig2^Cre^; R26-DTA^LSL^* (right) and controls (left) immunostained for CD31 (grey). Insets show the vasculature from inner (blue boxes) and outer (orange boxes) adrenal regions. **(N)** Schematic of late MN ablation by DT intraamniotic injection in *Olig2^Cre^; R26-iDTR^LSL^* embryos and control littermates. DT administration at E13 enables efficient adrenal gland denervation after SCP placement and differentiation. **(O)** Transverse spinal cord sections of E16.5 control embryos (left) and *Olig2^Cre^; R26-iDTR^LSL^* (right) upon DT injection at E13. *MN^(218-2)^::GFP* transgene identifies all MNs; nNOS labels only visceral MNs (vMNs). Asterisks indicate MN loss in *Olig2^Cre^; R26-iDTR^LSL^*. **(P)** Adrenal gland sections from E16.5 *Olig2^Cre^; R26-iDTR^LSL^*(right) and control (left) embryos. Visceral motor axon terminals are identified by *MN^(218-2)^::GFP* signal (green); CD31 labels endothelial cells (red); TH (blue) identifies chromaffin cells. The medulla is outlined. (**Q**) nNOS^+^ cell area within the vMN domain. (**R**) adrenal gland size. (**S**) TH^+^ adrenal chromaffin cell area. (**T**) CD31^+^ medullary vessel area in *Olig2^Cre^; R26-iDTR^LSL^*and controls. Scale bars: 100 µm. da: dorsal aorta; SCPs: Schwann cell precursors; sg: sympathetic ganglion.

Ablation of motor axons from the onset of adrenal gland innervation (E12) caused defects in vascular morphogenesis. To determine whether the influence of motor axons extends beyond this early developmental window, we employed an inducible ablation strategy. At E13, after adrenal innervation and SCP migration were largely complete (Akkuratova *et al*, 2022), Diphtheria toxin (DT) was administered via *in utero* intraamniotic injection (Fukunaga *et al*, 2018) to *Olig2^Cre^; R26-iDTR^LSL^* transgenic embryos expressing the human Diphtheria Toxin Receptor in motor neurons (Figure 2N). By E16.5, these embryos exhibited a near-complete loss of motor neurons in the spinal cord (Figure 2O, Q) and a corresponding lack of adrenal innervation (Figure 2P). Consistent with axon-mediated SCP placement being completed prior to motor neuron ablation, both adrenal gland size and medullary area remained unchanged (Figure 2P, R, S), and the organization of the adrenal vascular network was likewise preserved (Figure 2P, T).

These findings define an early developmental window during which visceral motor axons regulate adrenal organogenesis by controlling SCP positioning and vascular patterning.

### Semaphorin-3C from motor axons and SCPs restricts angiogenesis and regionalizes the adrenal vasculature

Our results so far show a spatiotemporal overlap between innervation and vascular remodeling in the developing adrenal gland. Loss of motor axons —and consequently depletion of SCPs/chromaffin cells— leads to an uncontrolled expansion of a cortex-like vascular network. This suggests that motor axons or medullary cells may secrete anti-angiogenic signals that ensure proportionate vascular growth. To dissect the pathways underlying adrenal vascular patterning, we integrated single-cell transcriptomic datasets from the adrenal gland (E13.5 to P5; (Hanemaaijer *et al*, 2021), medullary neural crest progeny (E12.5-E13.5; (Furlan *et al*, 2017) and motor neurons (E12.5; (Amin *et al*, 2021) (Figure 3A-B), and used CellChat software to infer communication pathways among adrenal cell populations (Figure 3C). When focusing on pathways involved in vascular patterning (Figure 3D), stromal cells (primarily derived from the external fibrous capsule) (Hanemaaijer *et al*, 2021) and cortical cells were predicted to signal prominently to ECs by releasing the pro-angiogenic factor VEGF (Figure 3E), while preganglionic (visceral) motor neurons (PGCa, PGCb), and to a lesser extent SCPs, expressed class-3 Semaphorins (SEMA3) known to restrict angiogenesis by eliciting endothelial cell repulsion (Martins *et al*, 2022; Vieira *et al*, 2022) (Figure 3F). In contrast, chromaffin cells showed limited signaling toward endothelial cells (Figure 3F). This computational analysis suggests a different regulation of vascular patterning in the two adrenal compartments, with the cortex displaying a greater angiogenic potential than the medulla. Both VEGF and SEMA3 signaling were among the top-ranked predicted cell interaction pathways in the developing adrenal gland (Figure 3C). When ligand-receptor pairs mediating cell repulsion were resolved, the strongest signal involved Semaphorin-3C (Sema3C) produced by motor neurons and SCPs, acting on endothelial cells through Plexin-D1 and Neuropilins (Nrp1, Nrp2) (Figure 3G). Previous studies have shown that Plexin-D1 mediates Sema3C signaling via Nrp1/2 co-receptors to induce endothelial repulsion, counterbalancing VEGF/VEGFR2 activity in other developmental contexts (Martins *et al*, 2022; Vieira *et al*, 2022; Plein *et al*, 2015). Supporting these predictions, scRNA-seq data showed that *Sema3C* expression was restricted to motor neuron and SCP clusters, whereas *Vegfa* was enriched in cortical and stromal cells (Figure S3A, B).

**Figure 3:**
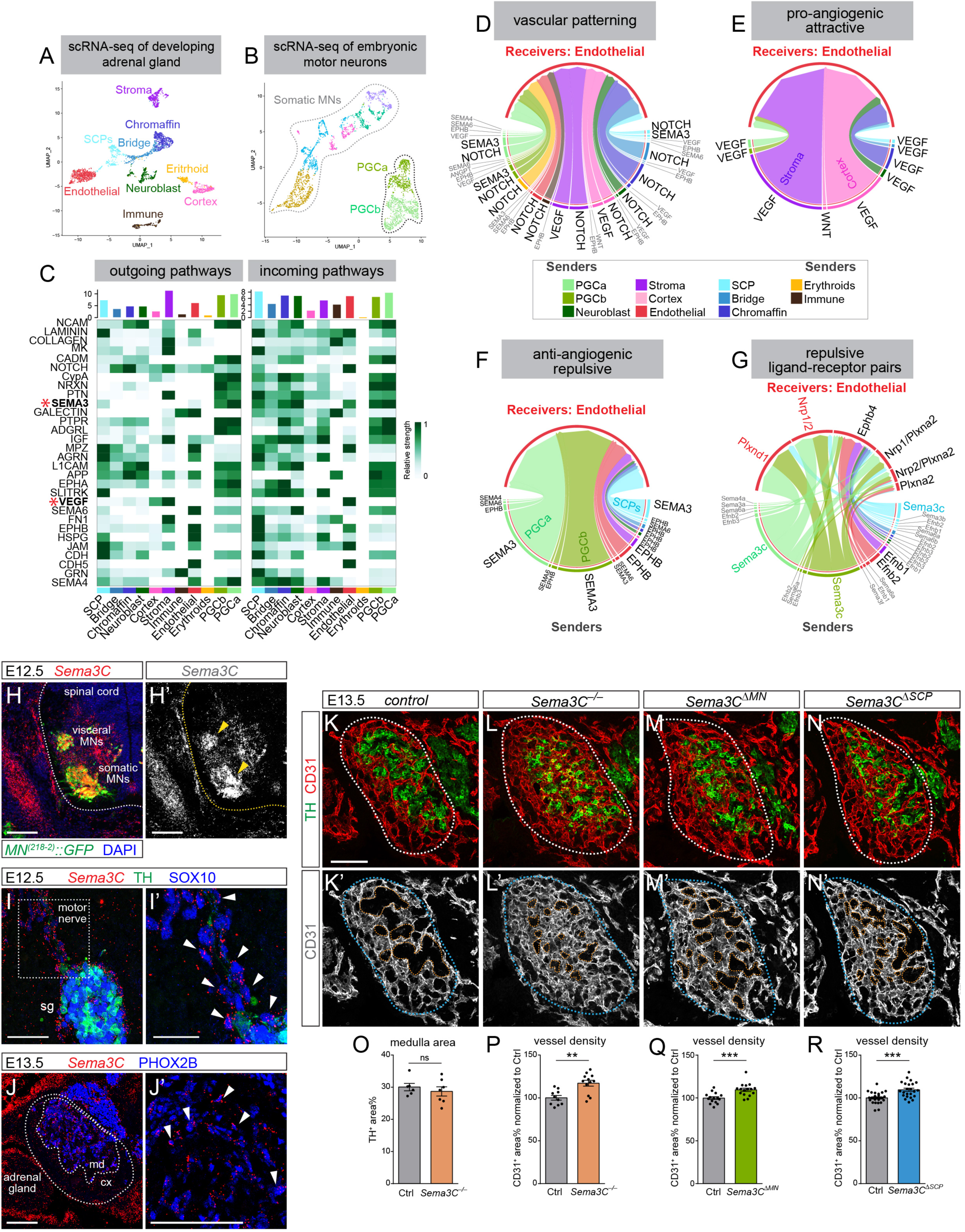
Sema3C signaling regulates medullary angiogenesis. **(A)** Integrated UMAP of scRNA-seq datasets of the developing adrenal gland from Hanemaijer et al 2021 and Furlan et al 2017. The main cell type clusters are labelled. **(B)** UMAP of scRNA-seq dataset of embryonic MNs from Amin et al 2021. Visceral MN clusters are labelled as PGCa and PGCb (preganglionic column). **(C)** Top 30 outgoing (right) and incoming (right) pathways mediating interactions among adrenal gland cell pairs. Color intensity identifies the strength of cell-cell communication. SEMA3 and VEGF pathways are highlighted (red asterisks). **(D – G)** Chord diagrams showing the strength of interactions between all adrenal gland cell types (senders) vs. endothelial cells (receivers) from curated lists of vascular patterning pathways (D), pro-angiogenic/attractive pathways (E), and pro-angiogenic/repulsive pathways (F). Ligand-receptor pairs mediating EC repulsion are shown in G. **(H – J’)** *Sema3C* (red) detected by RNAscope in spinal motor neurons (*MN^(218-2)^::GFP*^+^, green) at E12.5 (H, H’, arrowheads point to motor columns); migrating SCPs (SOX10, blue) at E12.5 (I, I’, arrowheads; boxed area in I is magnified in I’); adrenal SCPs (PHOX2B^+^, blue) at E13.5 (J, J’, arrowheads; J’ is magnified from J). **(K – N’)** Adrenal gland sections from E13.5 controls (K), *Sema3C -/-* (L), *Olig2^Cre^; Sema3C ^Flox/-^* (*Sema3C ^ΔMN^*) (M) and *Plp^CreER^; Sema3C ^Flox/-^* (*Sema3C ^ΔSCP^*) (N). TH (green) identifies chromaffin cells, CD31 (red) vessels. CD31 is shown separately (grey) in K’, L’, M’, N’. The adrenal gland is outlined. Medullary compartments are outlined in orange. **(O)** Area occupied by chromaffin cells in the adrenal gland of *Sema3C^−/−^* and controls. **(P – R)** Vessel density in the adrenal medulla of *Sema3C^−/−^* (P), *Sema3C^ΔMN^* (Q) and *Sema3C^ΔSCP^*(R) vs. control littermates. Scale bars: H, H’, 100 µm; I, I’, 50 and 25 µm; J, J’, 100 µm; K – N’, 100 µm.

In agreement with the transcriptomic analysis, RNAscope *in situ* hybridization detected high levels of *Sema3C* in visceral motor neurons within the spinal cord (Figure 3H, H’), as well as along their nerve tracts, where a more discrete signal was observed in migrating SCPs (SOX10^+^) (Figure 3I, I’) and in adrenal gland–resident SCPs (PHOX2B^+^) (Figure 3J, J’). Given that Sema3C signaling regulates spinal cord vascularization and motor axon-vessel interactions (Martins *et al*, 2022; Vieira *et al*, 2022), we investigated its contribution to adrenal angiogenesis. Specifically, we asked whether loss of Sema3C could recapitulate the vascular phenotype observed in *Olig2^Cre^; R26-DTA^LSL^* embryos, in which motor neurons and SCPs —the main sources of Sema3C during adrenal morphogenesis— are depleted. Unlike motor neuron-ablated embryos, *Sema3C* knockout embryos (*Sema3C^−/−^*) displayed normal medulla size (Figure 3K-L, O) but vessel density was notably increased in the medulla, blurring the typical cortical-medullary vascular zonation (Figure 3K, L’, P). This phenotype was penetrant, observed in ∼75% of E13.5 Sema3C^−/−^ embryos, but never in heterozygous or wild-type littermates. These results suggest that Sema3C limits vascular expansion during adrenal development, supporting the segregation of cortical and medullary vascular domains.

To dissect the cellular sources of this anti-angiogenic pathway, we used a floxed *Sema3C* allele to generate conditional knockout models, enabling selective deletion in either motor neurons (using *Olig2^Cre^*) or SCPs (using *Plp^CreER^* activated by tamoxifen at E10.5 and E11.5) (Figure S3H-I’). RNAscope at E13.5 confirmed targeted depletion of *Sema3C* in adrenal SCPs of *Plp^CreER^; Sema3^Flox/−^* embryos (hereafter, *Sema3C^ΔSCP^*) (Figure S3L-M’; compare with *Sema3C^−/−^*, Figure S3J-K’) and in motor neurons in *Olig2^Cre^; Sema3^Flox/−^* embryos (hereafter, *Sema3C^ΔMN^*) (Figure S3N-O’). In both conditional mutants, the adrenal gland exhibited abnormalities resembling those seen in full knockouts, though with reduced severity and penetrance (Figure 3M-N’, Q, R). These vascular defects were not observed in motor neuron-specific knockouts of pro-angiogenic ligands, such as *Vegfa* and Angiopoietin1 (*Ang1*) (Figure S3B-G), supporting the specificity of motor-derived anti-angiogenic Sema3C in regulating adrenal vascular patterning.

Together, these findings indicate that Sema3C derived from both motor axon terminals and their associated SCPs contributes to the local control of angiogenesis in the adrenal medulla, acting through spatially coordinated sources during gland colonization and innervation.

### Plexin-D1 is required in endothelial cells to coordinate adrenal vascular regionalization and medullary morphogenesis

Sema3C signaling through the endothelial receptor Plexin-D1 has been implicated in physiological angiogenesis in the spinal cord (Martins *et al*, 2022; Vieira *et al*, 2022) and was also identified *in silico* as a key pathway mediating cell–cell communication in the adrenal gland (Figure 3G). PlexinD1 is broadly expressed by endothelial cells throughout the developing embryo, including the adrenal vasculature (Figure S4A, A’). We therefore investigated the involvement of Plexin-D1 in adrenal vascular patterning by analyzing knockout embryos for this receptor. As early as E12.5, *Plxnd1^−/−^* embryos exhibited increased vascular density in the adrenal gland (Figure I), which became more pronounced by E13.5 (Figure 4C-D’, I). Despite these vascular changes, overall adrenal gland size remained unaffected (Figure 4J), with only a modest reduction in medullary cell number (∼10–20%) (Figure 4K). Notably, similar to *Sema3C* mutants, the adrenal vasculature lost its typical regional specificity in *Plxnd1^−/−^* embryos, with medullary vessels adopting a more uniform network topology resembling that of the cortical vasculature (Figure 4D, D’). To determine whether these alterations stemmed from Plexin-D1 activity in endothelial cells, we crossed a floxed *Plxnd1* allele to the endothelial-specific *Cdh5^CreER^* line and induced gene deletion with three consecutive daily injections of tamoxifen starting at E10.5 (Figure 4A and Figure S4A’, B). While the adrenal medulla size in E13.5 *Cdh5^CreER^; Plxnd1^Flox/Flox^* embryos (hereafter *Plxnd1^ΔEC^*) was comparable to controls (Figure 4M), vessel density increased (Figure 4E-F’, L), and this phenotype persisted at E15.5 (Figure S4E-G).

**Figure 4:**
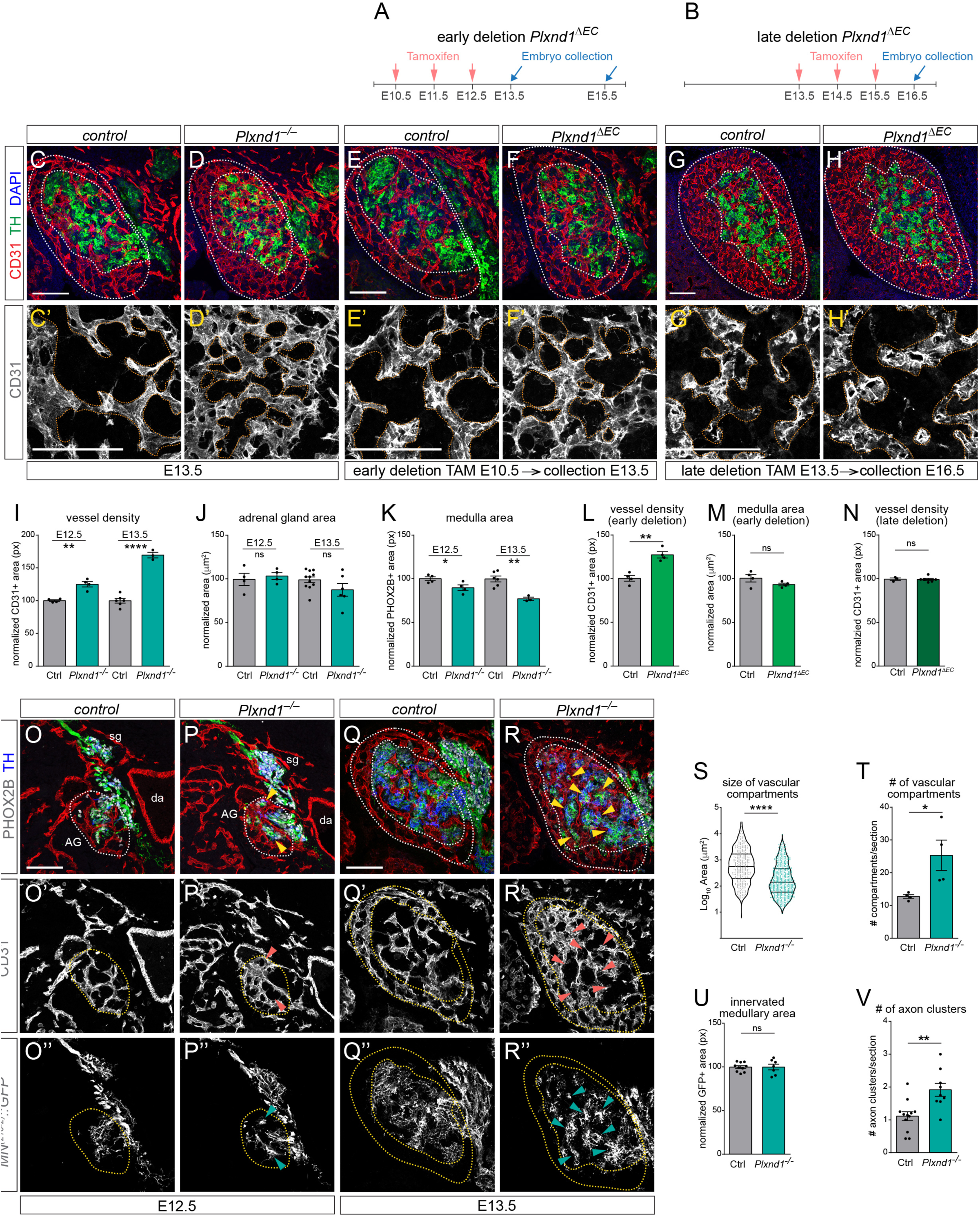
Plexin-D1 controls vascular patterning in the adrenal gland in a defined developmental window. **(A – B)** Timing of tamoxifen administration and embryo collection for early (A) and late (B) endothelial *Plxnd1* deletion in *Plxnd1 ^ΔEC^* embryos. **(C – H’)** Adrenal glands sections from E13.5 *Plxnd1^−/−^* (D, D’) and controls (C, C’); early-excised *Plxnd1 ^ΔEC^* embryos (F – F’, E13.5) and controls (E, E’); late-excised *Plxnd1 ^ΔEC^* embryos (H – H’, E16.5) and controls (G, G’). Immunostaining for TH (green) and CD31 (red). Nuclei (DAPI) are in blue. C’-H’ show a magnified view of adrenal vessels (CD31, grey) from the corresponding top panels. Medullary compartments are outlined in orange. **(I – K)** Adrenal (E12.5) and medullary (E13.5) vessel density (I), adrenal gland size (J), number of medullary cells identified as the area occupied by PHOX2B^+^ cells (K) in E12.5 and E13.5 *Plxnd1^−/−^* and controls. **(L, M)** Adrenal vessel density (L) and adrenal medulla area (M) in E13.5 controls and *Plxnd1 ^ΔEC^* upon early deletion of *Plxnd1*. **(N)** Vessel density in the adrenal gland of E16.5 controls and *Plxnd1 ^ΔEC^*upon late deletion of *Plxnd1*. **(O – R’’)** Transverse sections of E12.5 (O – P’’) and E13.5 (Q – R’’) *Plxnd1^−/−^* (P, R) and control (O, Q) embryos. The adrenal gland and medulla are outlined. Adrenal innervation is identified with *MN^(218-2)^::GFP* (green), mature and immature chromaffin cells are labelled with TH (blue) and PHOX2B (grey), respectively; vessels are marked with CD31 (red). CD31 and GFP channels are shown separately in grey in O’, P’, Q’, R’ and O’’, P’’, Q’’, R’’, respectively. Arrowheads point to aberrant visceral motor axon clustering. **(S)** Size distribution of medullary compartments defined by vascular plexus. Dots correspond to individual compartments. **(T)** Number of medullary compartments identified by the vascular plexus per section in *Plxnd1^−/−^* and controls at E13.5. **(U)** Area occupied by *MN^(218-2)^::GFP*^+^ visceral motor axons in the adrenal medulla of *Plxnd1^−/−^* compared to controls at E12.5 and E13.5. **(V)** Number of medullary axon clusters (identified by arrowheads in P’’ and R’’) per section in *Plxnd1^−/−^* and controls at E12.5 and E13.5. Scale bars: 100 µm. AG, adrenal gland; da, dorsal aorta; sg: sympathetic ganglion.

In contrast, late deletion of *Plxnd1* with tamoxifen injections starting at E13.5 (Figure 4B and Figure S4C-D) did not disrupt adrenal vascular patterning in *Plxnd1^ΔEC^* embryos (Figure 4G-H’, N). These experiments identified an early developmental window —coinciding with the entry of motor nerves and SCPs into the adrenal primordium— during which Plexin-D1 is required in endothelial cells to establish proper vascular architecture in the adrenal gland.

The adrenal vascular phenotypes of *Plxnd1^−/−^* embryos were more pronounced than those observed in *Sema3C* mutants, possibly reflecting the involvement of additional Semaphorin ligands that signal through Plexin-D1 (e.g., Sema3E, Sema4A, Sema3G) (Gu *et al*, 2005; Toyofuku *et al*, 2007; Liu *et al*, 2016). However, unlike *Sema3C* knockout, deletion of the canonical Plexin-D1 ligand Sema3E did not cause excessive adrenal vessel growth (Figure S4I-K), consistent with its minimal expression in both the developing adrenal gland and motor neurons (Figure S4H). Moreover, compound *Sema3C*/*Sema3E* knockout embryos displayed an increase in vessel density comparable to that seen in single *Sema3C* mutants (Figure S4I, L, M), further supporting Sema3C as a primary mediator of Plexin-D1 signaling during adrenal gland vascularization.

In *Plexin-D1* mutants, aberrant vascular expansion within the medulla disrupted the spatial organization of medullary components (Figure 4O-R”). The denser, more disorganized vascular network partitioned the medulla into a greater number of smaller compartments compared to controls, effectively confining chromaffin cells within these reduced spaces (Figure 4P, R, S, T). Furthermore, tracing visceral motor projections with the *MN^(218-2)^::GFP* revealed abnormal grouping of axon terminals in the medulla (Figure 4O”-R”). Whereas in controls motor axons were evenly distributed to spatially match chromaffin units, in *Plexin-D1* mutants they became aberrantly clustered, resulting in uneven innervation (Figure 4P”, R”, V). This disrupted spatial pattern suggests that the precise targeting and distribution of visceral motor axons is compromised, likely due to the altered vascular landscape and medullary architecture. Despite these changes, the overall thickness of the white ramus formed by visceral motor axons and the number of motor fibers reaching the medulla appeared preserved in *Plxnd1^−/−^* embryos (Figure 4U and Figure S4N, O). However, some visceral motor axons in *Plxnd1^−/−^* mutants may have been misrouted along their projection paths due to aberrant interactions with vessels, as shown for axial nerves (Martins *et al*, 2022).

These results demonstrate that adrenal endothelial cells respond to repulsive Sema3C signals from medullary components —motor axons and SCPs— via the Plexin-D1 receptor. Loss of endothelial Plexin-D1 leads to ectopic vascular expansion within the medulla, disrupting the normal regionalization of cortical and medullary vascular networks. This altered vascular architecture perturbs spatial relationships among blood vessels, chromaffin cell clusters, and visceral motor axons, compromising the precise cellular organization required for proper adrenal morphogenesis.

### Sema3C/Plexin-D1 signaling curbs a feedback loop of vascular mispatterning and adrenal cortical overgrowth

Although cortex and medulla develop interdependently (Bechmann *et al*, 2021; Kastriti *et al*, 2020), they exhibit distinct vascular patterns, suggesting differential affinities for the vasculature. Consistent with this, *in silico* cell-cell communication analysis predicted a predominance of pro-angiogenic signaling in cortical cells, whereas medullary components expressed anti-angiogenic and repulsive cues (Figure 3E, F). Conversely, endothelial cells were predicted to repel chromaffin —but not cortical— cells, primarily via Ephrin-A/B and Semaphorin signaling (Figure S5A). Reflecting these compartment-specific interactions, we found that vascular expansion —triggered by either genetic ablation of visceral motor nerves (*Olig2^Cre^; R26-DTA^LSL^*) or inactivation of Plexin-D1 signaling (*Plxnd1^−/−^*)— was accompanied by a striking overgrowth of the adrenal cortex (Figure 5A-N). In both models, SF1^+^ cortical cells spread abnormally into medullary territory, leading to intermingling of the two compartments (Figure 5 G”, K”). This was particularly evident in Plexin-D1 mutants, where the medulla was still present (Figure 4D, K), in contrast to its near-complete loss in *Olig2^Cre^; R26-DTA^LSL^* embryos (Figure 2B). Adrenal cortical enlargement was paralleled by a marked upregulation of *Nr5a1*, the gene encoding for SF1, which drives adrenocortical cell proliferation (Doghman *et al*, 2007). Notably, SF1^+^ cortical cells did not scatter throughout the medulla randomly. Instead, their migration closely followed the path of the ectopically expanded vasculature. Together, these findings indicate that in the absence of medullary-derived repulsive signals —such as Sema3C from motor axons and SCPs— vascular patterning becomes dysregulated, allowing cortical cells to exploit the aberrant vasculature as a scaffold to invade medullary territory, ultimately disrupting the spatial compartmentalization of the developing adrenal gland.

**Figure 5:**
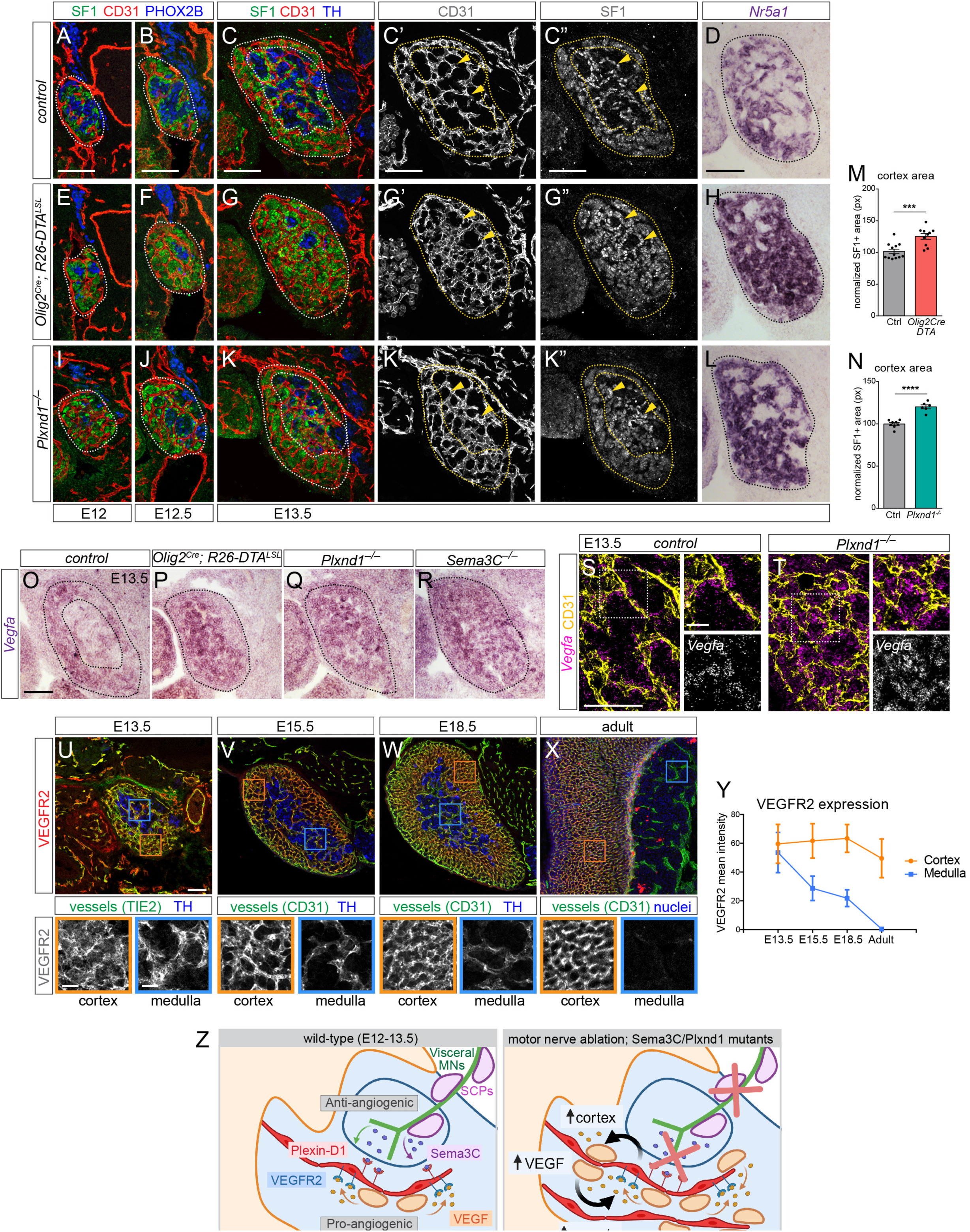
Motor innervation and PlexinD1-Sema3C signaling regulate proportionate expansion of the adrenal cortex. **(A – L)** Time-course of adrenal cortex development in E12 (A, E, I), E12.5 (B, F, J) and E13.5 (C, G, K) controls (A – C’’), *Olig2^Cre^; R26-DTA^LSL^* (E – G’’), *Plxnd1^−/−^* (I – K’’). Adrenal cortex (SF1^+^, green) and medulla (either PHOX2B^+^ or TH^+^, blue) are outlined. CD31 marks the adrenal vessels (red). C’, G’, K’ and C’’, G’’, K’’ show separate channels from E13.5 sections for CD31 and SF1, respectively (grey). Yellow arrowheads show association of cortical cells with medullary vessels. D, H, L show expression of SF1-encoding gene *Nr5a1* identified through *in situ* hybridization in E13.5 controls (D), *Olig2^Cre^; R26-DTA^LSL^* (H) and *Plxnd1^−/−^* (L). **(M – N)** Adrenal cortex size identified as SF1^+^ area within the adrenal gland of *Olig2^Cre^; R26-DTA^LSL^* (M) and *Plxnd1^−/−^* (N) versus the relative control littermates at E12.5 and E13.5. **(O – R)** *In situ* hybridization on adrenal sections from E13.5 control (O), *Olig2^Cre^; R26-DTA^LSL^* (P), *Plxnd1^−/−^* (Q) and *Sema3C^−/−^*(R) shows expansion of *Vegfa* expression domain in mutants. **(S – T)** Insets of adrenal gland sections from Figure S5A showing *Vegf* expression (identified by RNAscope, magenta), distributing along ECs (CD31^+^, yellow) in controls (S) and *Plxnd1^−/−^* (T) at E13.5. Right insets show a magnification of the boxed area. *Vegf* expression, shown separately in grey, is expanded within the medulla in *Plxnd1-/-* compared to controls. **(U – X)** Time-course of VEGFR2 expression (red) in adrenal vessels (identified with either CD31 or TIE2, green) at E13.5 (U), E15.5 (V), E18.5 (W) and postnatal stages (X). TH identify the medulla, DAPI stains cell nuclei. Insets with VEGFR2 expression in grey show a magnification of the boxed area; blue boxes identify medullary vessels, orange boxes identify cortical vessels. (**Y)** Quantification of VEGFR2 expression levels identified as mean signal intensity within cortical (orange) and medullary (blue) regions in adrenal gland sections of E13.5, E15.5, E18.5 and adult wild type mice. **(Z)** Schematic showing balanced regulation of adrenal angiogenesis and compartment formation in controls (left) vs how these processes go awry upon motor ablation or disruption of Sema3C-Plxnd1 signaling (right). Anti-angiogenic cues from the medullary components counteract pro-angiogenic signals from the cortex to ensure proportionate vessel growth in the medulla. When communication between axons-SCPs-vessels is disrupted, vessel overgrowth, due to loss of angiogenic inhibition, leads to dramatic expansion of cortical cells and increased VEGF release, which further sustains adrenal angiogenesis, thus establishing a detrimental vicious cycle. Scake bars: A – L, 100 µm. O – R, 100 µm; S – T, 100 µm; magnified insets in S – T, 25 µm; U – X, 100 µm.

Cortical cells were expected to stimulate angiogenesis through VEGF secretion (Figure 3E). Accordingly, *Vegfa* levels were significantly higher in the adrenal cortex than in the medulla of E13.5 control embryos (Figure 5O). In *Olig2^Cre^; R26-DTA^LSL^*, *Plxnd1^−/−^*, *Sema3C^−/−^*mutants cortical expansion into medullary territory was accompanied by a corresponding broadening of the *Vegfa* expression domain (Figure 5P-R). In control embryos, *Vegfa*^+^ cortical cells ensheathed vessels, and remained closely associated with them as the vasculature expanded abnormally across the medulla in *Plxnd1^−/−^* mutants (Figure 5S-T and Figure S5B). Consequently, the redistribution of Vegfa⁺ cortical cells into the medulla led to a loss of VEGF compartmentalization, resulting in a more uniform expression throughout the adrenal gland (Fan *et al*, 2024).

The differential cortex-high/medulla-low distribution of VEGF was mirrored by a developmental downregulation of its cognate receptor KDR/VEGFR2 in medullary vessels. At E13.5, VEGFR2 was expressed at comparable levels in both the cortex and medulla (Figure 5U). However, while cortical vessels maintained VEGFR2 expression into adulthood, VEGFR2 levels declined in medullary vessels after E15.5, becoming undetectable in the adult adrenal medulla (Figure 5V-Y). Therefore, during early stages of adrenal gland development, both medullary and cortical endothelial cells are competent to respond to pro-angiogenic VEGF signals produced by nearby vessel-associated cortical cells, which appear to be a major source of this ligand. Subsequently, however, medullary vessels undergo a rapid shift to a VEGFR2-low, and eventually VEGFR2-off, state that renders them progressively unresponsive to VEGF signaling.

Collectively, these findings support a model in which loss of adrenal innervation or inactivation of Sema3C/Plexin-D1 dampens anti-angiogenic signaling required to counterbalance VEGF/VEGFR2-driven vascularization. Consequently, vascular growth proceeds unchecked, leading to ectopic expansion of blood vessels into the medulla. This initiates a detrimental feedback loop —a vicious cycle— of vascular mispatterning and cortical overgrowth, as the expanding vasculature draws VEGF-secreting cortical cells into medullary territory, further fueling aberrant vessel growth and cortical invasion. As a result, the spatial compartmentalization of the adrenal gland is progressively lost. Thus, proportional and compartmentalized vascular development orchestrated by these antagonistic signals is critical for the structural integrity of the organ (Figure 5Z).

### Vascular mispatterning and cortical expansion promote aberrant macrophage-mediated tissue remodeling

Tissue overgrowth and architectural disruption during fetal development can lead to cellular stress and sustained damage (Lawrence *et al*, 2024). To investigate the impact of cortical expansion linked to vascular mispatterning on adrenal organogenesis, we examined mutant embryos at late gestational stages —the latest timepoints available given the perinatal lethality of our genetic models. By E16.5, *Olig2^Cre^; R26-DTA^LSL^* embryos and, although to a lesser extent, *Plxnd1^−/−^*embryos, showed a striking accumulation of apoptotic cells, identified by Caspase-3 staining, specifically at the cortex–medulla boundary (Figure 6A-C’, G, H). This localized apoptotic burst was accompanied by the activation of an innate inflammatory response, marked by accumulation of IBA^+^ macrophages (Figure 6D-F’, I, J). An increased number of macrophages was already detectable at E15.5 in *Olig2^Cre^; R26-DTA^LSL^*, *Plxnd1^−/−^,* and *Sema3C^−/−^* embryos, with varying degrees of severity (Figure S6A-D”). While macrophage accumulation was particularly evident at the cortex–medulla transition zone, it often extended beyond the region of apoptotic cell death (Figure 6E’, F’). In *Olig2^Cre^; R26-DTA^LSL^* and *Plxnd1^−/−^*embryos, these macrophages exhibited a globular morphology characteristic of an activated, phagocytic state and were observed engulfing apoptotic cell debris labeled by TUNEL staining (Figure 6K-M and Figure S6E-F”). Consistent with this, they displayed a CD206^+/^CD86^low^ anti-inflammatory phenotype, typical of phagocytic macrophages engaged during physiological apoptotic events in embryonic development (Gregory & Devitt, 2004; Weavers *et al*, 2016) (Figure S6I-J’’’). By E18.5, apoptotic cells were largely cleared, yet macrophages with an activated morphology persisted at these later stages (Figure S6G-H’). The cortex– medulla interface where apoptotic cells, and hence phagocytic macrophages, prominently accumulated in mutant embryos, is a site of adrenocortical turnover during normal postnatal maturation (Chang *et al*, 2011, 2013; Carsia *et al*, 1996). Accordingly, apoptotic and phagocytic events were observed in wild-type mice around puberty (3-4 weeks of age) at the level of regressing fetal cortex (X-zone), suggesting that apoptotic cortical cells are normally cleared by innate immune cells as part of the physiological postnatal remodeling of the adrenal gland (Figure 6N-S).

**Figure 6:**
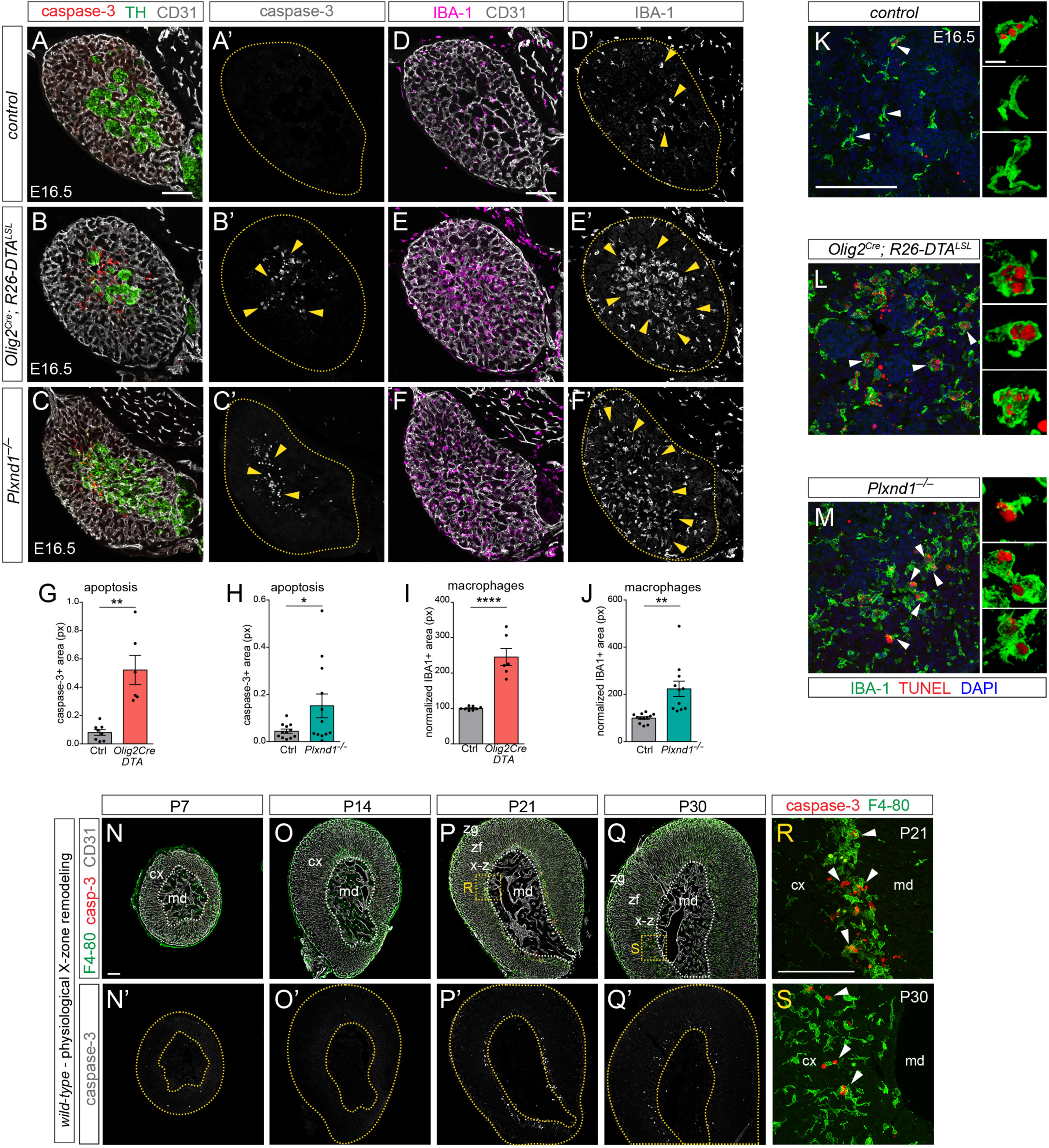
Aberrant phagocytic macrophage response induced by vascular mispatterning and cortical expansion. **(A – C’)** Adrenal gland sections of E16.5 controls (A, A’), *Olig2^Cre^; R26-DTA^LSL^* (B, B’) and *Plxnd1^−/−^* (C, C’) immunostained for CD31 (grey), TH (green) and active (red) which labels apoptotic cells. A’, B’, C’ shows apoptosis (caspase-3 signal, arrowheads) in grey. The adrenal gland is outlined. **(D – F’)** E16.5 adrenal sections of controls (D, D’), *Olig2^Cre^; R26-DTA^LSL^*(E, E’) and *Plxnd1^−/−^* (F, F’) immunostained for CD31 (grey) and macrophage marker IBA1 (magenta). D’, E’, F’ shows macrophage accumulation (grey, arrowheads) in mutants (E’, F’) but not controls (D’). The adrenal gland is outlined. **(G – J)** Area of caspase-3^+^ signal (G, H) and IBA1^+^ macrophages (I, J) within the adrenal gland of *Olig2^Cre^; R26-DTA^LSL^* (G, I) and *Plxnd1^−/−^* embryos (H, J) compared to controls. **(K – M)** Medullary insets showing phagocytic macrophage activation in *Olig2^Cre^; R26-DTA^LSL^* (L) and *Plxnd1^−/−^*(M) but not in control embryos (K) at E16.5. IBA1^+^ macrophages are in green; apoptotic cell debris are identified by TUNEL (red). DAPI (blue) identifies nuclei. Arrows point to quiescent macrophages with elongated morphology; arrowheads identify examples of activated phagocytic macrophages with globular morphology. Insets show magnified vies of representative macrophage morphologies. **(N – S)** Adrenal sections from post-natal wild-type male mice from P7 to P30 immunostained for caspase-3 (red), CD31 (grey) and the macrophage marker F4-80 (green). Caspase-3 staining is shown separately (N’ – Q’). Cortex and medulla are outlined. R and S show a magnified view of apoptotic cells (red) and macrophages (green) from the boxed area in P and Q, respectively. cx: cortex; md: medulla; x-z: x-zone; zf: zona fasciculata; zg: zona glomerulosa. Scale bars: 100 µm; magnified panels in K – M, 10 µm.

We conclude that early disruption of neurovascular patterning compromises adrenal compartmentalization and drives cortical overgrowth, triggering an exaggerated innate immune response that hijacks and amplifies a physiological program of cortical cell turnover.

## Discussion

A long-standing question in developmental biology is how distinct cell populations become organized into spatially and functionally segregated (Krens & Heisenberg, 2011; Salazar-Ciudad *et al*, 2003). Compartmentalization is a defining feature of organ architecture, allowing specialized cell types to localize within specific regions, interact with appropriate cues, and contribute to integrated tissue function (Fagotto, 2014). These spatial arrangements are not exclusive to embryonic development; they are also critical during adult tissue repair, where re-establishing boundaries is key to restoring structure and function. Conversely, the loss of tissue compartmentalization is a common feature of pathological conditions, including chronic inflammation, fibrosis, and cancer. The formation of such boundaries is traditionally attributed to cell-intrinsic programs and intercellular signaling mechanisms that mediate differential affinities between homotypic and heterotypic cell types—often through matching chemical labels such as cadherins, ephrins, or other surface receptors affinities (Fagotto *et al*, 2013; Taylor *et al*, 2017; Steinberg & Gilbert, 2004). In parallel, biomechanical features such as differential adhesion, cortical tension, migratory forces, and intercellular stress patterns also contribute to cell sorting and positioning (Lucia *et al*, 2022). Here, we reveal a distinct mechanism for cell segregation and boundary formation, in which the spatial architecture of the vasculature —not just the intrinsic chemical and physical properties of parenchymal cells— guides tissue partitioning during organogenesis. In the developing adrenal gland, cortical and medullary cells, which originate from different embryonic lineages and carry divergent signaling identities, do not segregate through direct repulsion or mechanical competition alone (Bornstein *et al*, 1994; Bechmann *et al*, 2021; Haase *et al*, 2010). Instead, their organization is orchestrated by a zonated vascular scaffold shaped by localized neurovascular interactions. Specifically, visceral motor axons and their associated SCPs deliver the repulsive, anti-angiogenic cue Sema3C, which signals through endothelial PlexinD1 to restrict vascular growth within the medulla. In parallel, cortical cells secrete pro-angiogenic VEGF, driving vessel expansion in the surrounding cortex. This antagonistic interplay gives rise to two spatially distinct vascular microenvironments: a dense, highly vascularized cortex that supports the migration and expansion of cortical endocrine cells, and a sparsely vascularized medulla, where chromaffin cell clusters remain confined within a capillary network shaped by neurovascular signaling. This spatial organization reflects a differential affinity for the local vasculature. Cortical cells avidly associate with blood vessels, migrating along vascular tracks and remaining tightly apposed once settled in the cortex —consistent with their reliance on blood-borne cues for hormone release. Chromaffin cells, by contrast, localize near but not in direct contact with vessels —a proximity that, while enabling efficient hormone delivery into circulation, suggests active segregation from the vasculature. This is supported by the enrichment of adhesive ECM pathways mediating endothelial– cortical cell interactions, in contrast to the presence of potentially repulsive Semaphorin and Ephrin signaling between endothelial and chromaffin cells. We propose that differential vascular affinity, layered onto a pre-patterned neurovascular scaffold, drives the physical and functional partitioning of the adrenal gland. In this model, the vasculature does not simply adapt to surrounding tissues but actively imposes spatial order on the developing organ, guiding where specific cell types can localize, expand, and differentiate. Rather than arising from direct repulsion between cell types, compartment boundaries are shaped by differential access to vascular niches, which supports both proportional growth and functional segregation of endocrine compartments.

Neurovascular signaling establishes a self-reinforcing architecture: VEGF-producing cortical cells promote local angiogenesis and maintain vessel association, while Sema3C restricts vascular expansion into the medulla, preserving compartment boundaries. Disruption of this signaling balance—through loss of innervation or Semaphorin3C-PlexinD1 pathway ablation—leads to cortical cell invasion along ectopic vessels, excessive angiogenesis, and displacement of chromaffin cells. As boundaries blur and regional identity erodes, a macrophage response is triggered, reflecting maladaptive tissue remodeling that aberrantly recapitulates a physiological postnatal process of gland reorganization. Thus, spatial control of angiogenesis functions not only to deliver nutrients but to establish and maintain structural and functional compartmentalization during organogenesis. This adds a new layer to the emerging view of blood vessels as migratory substrates for progenitor cells or axons (Fujioka *et al*, 2019; Cattin *et al*, 2015; Tsai *et al*, 2016; Bhat *et al*, 2024). In the adrenal gland, however, the vasculature is pre-patterned by neurovascular signals to create a spatial template that serves as an instructive boundary—physically and functionally partitioning the tissue and helping define compartment identity. This work suggests that vascular partitioning may represent a conserved and versatile developmental strategy in organs that require tight spatial segregation of specialized cell populations. Similar antagonistic angiogenic guidance systems —such as the balance of VEGF and Semaphorin3C— have been described in the spinal cord, where motor neurons shape their own vascular territories to avoid interference with axon guidance (Martins *et al*, 2022; Vieira *et al*, 2022). This strategy may be particularly relevant to other neuroendocrine systems, and its disruption may underlie disease. For instance, loss of compartmental integrity and abnormal angiogenesis are hallmarks of adrenal tumors. Our findings raise the possibility that failure of vascular partitioning contributes to pathological cell mixing, aberrant signaling microenvironments, and ultimately neoplastic transformation. This may be particularly relevant in the adrenal gland, which is a common site of origin for neuroblastoma (Maris *et al*, 2007; Matthay *et al*, 2016). Elucidating how compartmental boundaries are established during normal development may thus offer insight into the mechanisms by which developmental programs are reactivated during tumor initiation and progression. Notably, tumor cell interactions with nerves and blood vessels have been shown to promote metastatic dissemination in neuroblastoma and other cancers (Delloye-Bourgeois *et al*, 2017; Ben Amar *et al*, 2022), highlighting the broader relevance of neurovascular signaling in both organogenesis and oncogenesis.

In summary, our findings reveal that vascular patterning serves as an instructive mechanism for organ compartmentalization. In the adrenal gland, neurovascular signals establish topographically organized vascular networks that guides the segregation of cortical and medullary cells according to their differential affinities for blood vessels. This strategy not only reinforces physical separation but also supports functional specialization, aligning spatial organization with endocrine output. More broadly, we propose that specialized vascular scaffolds coordinate local signaling with tissue architecture, and that their disruption may represent a common mechanism in developmental disorders and tumorigenesis.

## Acknowledgements

We thank Ueli Suter and Stefano Previtali for *Ppl-CreERT2* mice; Ralf Adams and Silvia Brunelli for *Cdh5(PAC)-CreERT2* mice; Jonathan Raper and Sophie Chauvet for *Sema3C^−/−^* mice; Genentech, Inc. for *Vegfa^flox^* mice; Samuel Pfaff and Neal Amin for *MN^(218-2)^::GFP* mice; Yutaka Yoshida for *Plxnd1^flox^* mice; Cesare Covino and the Advanced Microscopy Laboratory (ALEMBIC) of San Raffaele Hospital for expertise and instrumentation. Funding sources: Giovanni Armenise-Harvard Foundation Career Development Award and European Research Council Starting Grant 335590 to D.B. British Heart Foundation project grant PG/22/10935 to C.R.

## Author contributions

A.M. designed experiments, collected and analyzed data with the help of A.B., G.P.B., C.M., I.B., and C.S. S.D. performed computational analysis. A.S. performed in utero injections. E.I. and C.R. provided embryo samples and critical comments. D.B. supervised the study and wrote the manuscript with A.M.

## Materials and Methods

### Mouse lines

*Olig2::Cre* (Dessaud *et al*, 2007); *Cdh5(PAC)-CreERT2* (Wang *et al*, 2010); *Plp::CreERT2* (Leone *et al*, 2003); *MN^(218-2)^::GFP* based on MN-specific miR-218-2 promoter from the *Slit3* locus (Amin *et al*, 2015); *R26-Tomato^LSL^* (JAX stock# 007914) (Madisen *et al*, 2010); *Plxnd1*-null allele generated from crossing *Plxnd1^flox^*(JAX stock# 018319) (Zhang *et al*, 2009) with ubiquitous *EIIa::Cre* (JAX stock# 003724) (Lakso *et al*, 1996); *Sema3C^−/−^* (Feiner *et al*, 2001); *Sema3C^flox^* (Plein *et al*, 2015); *Sema3C*-null allele generated from *Sema3C^flox^* upon germline recombination (Cai *et al*, 2022) was combined with *Sema3E*^−/−^ (Gu *et al*, 2005) to generate double mutants; *Angpt1^flox^* (JAX stock# 28925) (Jeansson *et al*, 2011); *VEGF^flox^* (Gerber *et al*, 1999); *R26-DTA^LSL^* (JAX stock# 010527) (Wu *et al*, 2006); *R26-iDTR^LSL^* (Jax stock# 007900) (Buch et al, 2005).

Tamoxifen (Tmx; Sigma-Aldrich Cat# T5648) was dissolved at 10mg/ml in 1:9 ethanol:corn oil (Sigma-Aldrich Cat# S5007) and administered intraperitoneally (i.p.) to pregnant females at a dose of 2mg of Tmx per 30g of body weight. Conditional endothelial excision of *Plxnd1^flox^* with *Cdh5^CreER^* was induced upon three consecutive Tmx injections at E10.5, E11.5 and E12.5 (early deletion), or E13.5, E14.5 and E15.5 for late *Plxnd1* deletion. *Sema3C^flox^* deletion from SCPs with *Plp^CreER^* was induced by two consecutive Tmx injections at E10.5 and E11.5. For MN ablation by intra-amniotic injection of Diphtheria Toxin (DT) in *Olig2^Cr^*; *iDTR^LSL^* embryos, pregnant females were anesthetized by i.p. injection of Avertin (2,2,2-Tribromoethanol, 7.5mg per 30 g of body weight, Sigma) and the abdominal skin was sterilized. The uterus was exposed, and individual E13 embryos were intra-amniotically injected with 40ng of DT (Calbiochem #322326) diluted in 15µL of sterile water using a pulled glass microcapillary needle. Embryos were then collected at E16.5. DT stocks were prepared at 2mg/mL in DPBS under sterile conditions.

Embryos were generated by timed mating, and the day on of vaginal plug was designated as embryonic day 0.5 (E0.5). All alleles were maintained in C57/BL6N strain except: *Sema3C^−/−^*, *Plp^CreER^; Sema3C ^Flox/−^* and *Olig2^Cre^; Sema3C ^Flox/−^*mice, which were in CD1 strain; *Olig2^Cre^; R26-DTA^LSL^*in CB6F1 strain; for late gestation analysis (E16.5 - E18.5) *Plxnd1*-null allele was moved into CB6F1 background to reduce mutant embryo mortality, which occurred at ∼E15.5 in C57/BL6N strain. *Olig2^Cre^; iDTR^LSL^* males were crossed to wild-type CB6F1 females for intra-amniotic DT injection.

Mice were maintained in pathogen-free facilities under standard housing conditions with *ad libitum* access to food and water. All experimental procedures and animal handling were approved by the Animal Research Committee of IRCCS San Raffaele Hospital or at the Animal Welfare and Ethical Review Body at the Biological Resources Unit of the UCL Institiute of Opthalmology with permission from the UK Home Office, and performed in compliance with IACUC guidelines.

### Tissue preparation

Embryos were collected at different time points depending on the experimental design, from E11.5 up to E18.5, and fixed with 4% Paraformaldehyde (PFA) in PBS at 4°C from 3 to 7 hours depending on the embryo size. Embryos were washed in PBS, then cryoprotected with 30% sucrose in PBS for 4-8 hours at 4°C and embedded in OCT (Sakura 4583). Adrenal glands from post-natal wild type animals were fixed in 4% PFA in PBS at 4°C for 1.5-2 hours, cryopreserved in 30% sucrose in PBS at 4°C overnight (ON) and then embedded in OCT/Sucrose 1:1. Embryo and adrenal gland blocks were cryo-sectioned to obtain 20µm sections for both immunofluorescence/immunohistochemistry and *in-situ* hybridization/RNAscope experiments.

### Immunofluorescence on tissue sections

Air-dried slides were incubated with primary antibodies diluted in blocking solution (1% BSA, 0.2% TritonX-100 in PBS) ON at 4°C. The day after, embryo sections were incubated with secondary fluorescent antibodies diluted in blocking solution for 2 hours at room temperature (RT). Nuclei were stained with DAPI (Sigma D9542). Slides were mounted with Fluorescent Mounting Medium (Dako S3023).

*Primary antibodies used for immunostaining:* rabbit anti-Caspase3 (1:300; BD Pharm 559565); rabbit anti-CD206 (1:500; Abcam 64693); rat anti-CD31 (1:200; BD Biosciences 550274); goat anti-CD31 (1:1000; R&D systems AF3628); rat anti-CD86 (1:100; Biolegend 105008); rabbit anti-ChromograninA (1:500; Abcam ab254322); rabbit anti-Claudin5 (1:300 Invitrogen 34-1600); rat anti-F4-80 (1:300 Abcam ab6640); rabbit anti-GFP (1:5000; Thermo Fisher Scientific A6455); chicken anti-GFP (1:1000; Abcam ab13970); rabbit anti-IBA1 (1:500 Wako 019-19741); rabbit anti-nNOS (1:1000; Immustar 24287); goat anti-PDGFRβ (1:200; R&D systems AF1042); goat anti-PHOX2B (1:500; R&D systems AF4940); PLVAP (1:200 BD Bioscience 553849); rabbit anti-RFP (1:1000 MBL Life Science PM005);rabbit anti-SF1 (1:400; Proteintech 18658-1-AP); rabbit anti-SOX10 (1:1000; Abcam ab155279); rabbit anti-TH (1:1000; Millipore AB152); chicken anti-TH (1:500; Abcam ab76442); goat anti-Tie2 (1:200; R&D systems AF762-SP); rabbit anti-TRKA (); rat anti-vCAM1 (1:100 BD Pharm 550547); rabbit anti-VaChat (1:1000; Synaptic Systems 139103); rat anti-VEGFR2 (1:100; BD Pharm 550549). *Secondary antibodies used for immunostaining:* goat or donkey anti-goat/rabbit/rat Alexa Fluor 488-, 555-, 647-conjugated (1:1000; Thermo Fisher Scientific); donkey anti-chick Alexa Fluor-488 (1:500; JacksonImmunoResearch 703-545-155); donkey anti-rat Cy3 (1:500; JacksonImmunoResearch 712-165-153); donkey anti-rabbit Alexa Fluor-405 (1:250; JacksonImmunoResearch).

### Immunohistochemistry on tissue sections

Immunohistochemistry was carried out on 20µm cryostat sections. After washes in PBS, sections were dehydrated in an ascending series of methanol (25% MeOH in PBS; 50% MeOH in PBS; 75% MeOH in H_2_O; 100% MeOH), incubated in 3% H_2_O_2_ in MeOH for 15 minutes at RT to block endogenous peroxidases, and rehydrated in a descending methanol series. Sections were incubated in blocking solution (10% FBS −foetal bovine serum; 1% BSA; 0.1% TritonX-100 in PBS) for 1 hour at RT. Primary goat anti-PlexinD1(R&D system AF4160) was diluted 1:200 in blocking solution and incubated ON at 4°C. After washes in PBS, sections were incubated with secondary biotinylated rabbit anti-goat IgG (Vectastain elite ABC kit Vector lab PK-6105) diluted 1:200 in blocking solution for 1 hour and 30 minutes at RT. Immunoreactive signal amplification was obtained through incubation with ABC working solution (Vectastain elite ABC kit Vector lab PK-6105) for 1 hour and 30 minutes at RT. Signal detection was carried out using DAB HRP substrate kit (Vector lab SK-4100): sections were incubated with DAB (3,3’-diamiobenzidine tetrahydrochloride) staining solution for few minutes at RT (until the signal reaches the optimal intensity), then dipped in PBS to stop the reaction. Slides were mounted with Glycergel mounting medium (Dako C0563).

### In-situ hybridization and RNAscope

Embryo sections were post-fixed in 4% PFA in PBS for 20 minutes at RT and incubated with 0.5µg/mL Proteinase K (PK –Euroclone EMR023100) in 50mM Tris-HCl pH8, 5mM EDTA pH8, for 5 minutes at 30°C. The proteinase reaction was stopped with a quick wash in 0.2% Glycine in PBS and slides were fixed again with 4% PFA in PBS for 10 minutes at RT. Then, the sections were dipped in acetylation solution (acetic anhydride diluted 1:400 in 0.1M triethanolamine-HCl pH8) with constant stirring for 10 minutes. Slides were incubated in pre-hybridization solution (50% formamide; 3x SSC) for at least 2 hours at RT. Embryo sections were then incubated ON at 56-65°C with 1µg/mL digoxygenin (DIG)-labelled RNA probe diluted in hybridization solution (50% formamide, 3x SSC, 10% Dextran Sulphate, 0.5mg/mL yeast RNA [Sigma, R6750], 1x Denhardt’s reagent). The following day, slides were washed twice in 50% formamide, 2x SSC for 20 minutes at 56°C, then incubated with 20μg/mL RNAse A (Sigma R5000) in NTE (0.5M NaCl, 10mM Tris-HCl pH8, 5mM EDTA pH8) for 30 minutes at 37°C, and washed twice in 50% formamide, 1x SSC for 20 minutes at 56°C. After two washes in 2x SSC and two washes in TBS (100mM Tris-HCl pH7.5, 150mM NaCl), sections were blocked with 10% sheep serum in TBS for at least 1 hour at RT, then incubated with anti-DIG AP-conjugated antibody (Roche 11 093 274 910) diluted 1:2000 in blocking solution ON at 4°C. After 2 washes in TBS and 2 washes in NTM (100mM NaCl, 100mM Tris-HCl pH9.5, 50mM MgCl_2_), signal detection was performed incubating slides with staining solution (3µL NBT [Sigma 11383213001], 3µL BCIP [Sigma 11383221001] for each mL of NTM) in the dark at RT or at 37°C, until desired signal intensity was obtained. Colorimetric reaction was stopped with 3 washes in PBS and slides were mounted with Glycergel mounting medium (Dako, C0563).

*Antisense DIG-labelled RNA probes used for in-situ hybridization:* mouse *Vegfa* (GenBank: NM_001317041.1; 1431-1849bp) cloned in pGEM (Ruiz de Almodovar *et al*, 2011); mouse *Nr5a1* (GenBank: NM_139051.2; 1223-2164 bp).

RNAscope was performed using the Multiplex Fluorescent Reagent Kit v2 (ACD BioTechne 323100) according to the manufacturer’s instructions. *Probes used for RNAscope:* mouse *Sema3C* (ACD BioTechne 441441-C3); mouse *Vegfa* (ACD BioTechne 521471).

### Tunel assay

After running immunofluorescence staining, apoptotic cells were detected using the “*Apotag RED in situ apoptosis detection kit*” according to manufacturers’ instructions. Briefly, after 10 seconds incubation with *Equilibration Buffer* at RT, sections were incubated with *Working Strength TdT Enzyme* solution for 1 hour at 37°C, then reaction was stopped with *Working Strength STOP/WASH Buffer* for 10 minutes at RT. After three washes in PBS, anti-Digoxigenin-Rhodamine was applied for 30 minutes at RT. Nuclei were stained with DAPI (Sigma D9542). Slides were mounted with Fluorescent Mounting Medium (Dako S3023).

### Embryonic adrenal gland-HUVEC co-cultures

Adrenal glands from n=8-11 E14.5 wild-type embryos were manually dissected under a sterile dissection hood and placed into Eppendorf tubes with 300µL of dissociation buffer (1.8 mg/mL Collagenase type II [Gibco 17101015], 0.18 mg/mL DNAse I [Sigma 260913-25MU], 10% FBS [Euroclone ECS5000L], 1x PenStrep [Euroclone ECB3001D] in PBS with Ca^2+^ Mg^2+^). After incubation at 37°C for 20 minutes, samples were dissociated by pipetting up and down. Collagenase reaction was stopped by adding 1mL of blocking buffer (without Ca^2+^ Mg^2+^) on top. Adrenal gland cell suspensions were pelleted at 0.3 rcf for 5 minutes at RT and resuspended in 1mL of blocking buffer. Cell preparations were filtered using a 35µm cell strainer (Corning Falcon 352235) to remove tissue aggregates, then pelleted and resuspended in 250 µL of Adrenal Gland Medium (10% FBS, 1x PenStrep, 1x L-Glutamine [Euroclone ECB3000D] in DMEM/F12 [Thermo/Life Technologies 11039021] supplemented with 0.025 µg/mL basic-FGF [Miltenyi Biotec 130-104-922]. 90000 adrenal gland cells were plated together with 20000-30000 Human umbilical vascular ECs (HUVEC; Lonza C2519A) in EC Growth Basal Medium-2 (EBM-2; Lonza CC-3156) on 13mm nitric acid pre-treated glass coverslips coated with 100 µg/mL Poly-D-Lysine (Sigma P7280-5MG) and 5 µg/mL Laminin (Gibco Cat# 23017015). After 24-, 72- or 120-hours coverslips were fixed in 4% PFA / 4% sucrose for 25 minutes at RT. Immunocytochemistry was performed using anti-Phalloidin (F-actin) AlexaFluor-555 (1:1000 Thermo/LifeTech A34055), goat anti-human VE-Cadherin (1:100 R&D systems AF938-SP), chicken anti-TH (1:500; Abcam ab76442) primary antibodies in blocking solution (1% BSA, 0.1% TritonX-100 in PBS) for 2 hours and secondary antibodies (Alexa Fluor-conjugated 1:1000; Thermo Fisher Scientific) in blocking solution for 1 hour.

### Microscopy

Confocal images were acquired with an Olympus Fluoview FV3000RS or Leica TCS SP8 SMD FLIM Laser Scanning Confocal microscopes. Immunofluorescent images used for quantifications were acquired with Zeiss Axio Observer Z1 microscope. Immunohistochemistry and *in-situ* hybridization colour brightfield images were acquired with Zeiss Axio Imager M2m microscope.

### Image analysis and statistics

Images used for quantification were acquired with same microscope settings, adjusted to the same background levels whenever possible and processed with Fiji/ImageJ and Adobe Photoshop.

The size of the adrenal gland or primordium was measured by manually drawing the perimeter of the organ in each section based on DAPI staining, using the freehand selection tool in Fiji/ImageJ. Each area value in µm^2^ was normalized to the control mean value. Adrenal medulla area was calculated as the TH^+^ or PHOX2B^+^ pixel area normalized to the total adrenal area (area%) after applying a threshold in Fiji/ImageJ to generate a binary mask. The same procedure was used to quantify adrenal cortical area as SF1^+^ area%. All values were normalized to the mean of relative control littermates. Blood vessel density within the whole adrenal gland or medulla was measured as the CD31^+^ pixel area normalized to the total area of the gland/organ (area%) after applying a threshold in Fiji/ImageJ to generate a binary mask. The same procedure was applied to measure VEGFR2^+^ or Tie2^+^ vessel and PDGFRβ^+^ pericyte area within the adrenal gland. Quantification of the ratio between angiogenic and mature vasculature was calculated as the ratio between VEGFR2^+^ and Tie2^+^ vessel area%. Similarly, the pericyte coverage was calculated as the ratio between PDGFRβ^+^ area% and CD31^+^ vessel area%. All values were normalized to the mean of relative control littermates. The area of medullary compartments was measured by manually tracing the border of the empty spaces defined by the vascular network within the adrenal medulla. Each dot in Figure 2L and Figure 4S represents a single compartment; the size of each compartment in µm^2^ is expressed in Log_10_ units; the number of medullary compartments per section within the adrenal medulla area was plotted in Figure 2K and Figure 4T. The number of visceral motor neurons was identified by measuring the pixel area occupied by nNOS^+^ signal within an equal sized ROI drawn around the intermediolateral region of the spinal cord (Figure 2Q). For quantification of motor innervation defects in the adrenal medulla of *Plxnd1^−/−^,* the number of axon clusters (defined as GFP^+^ area > 20 pixel^2^) per section was manually counted (Figure 4V). To quantify the innervation of the adrenal medulla by visceral motor axons in, an equal threshold was applied to GFP staining. The area occupied by GFP^+^ pixels within the medulla was measured after generating a binary mask in Fiji/ImageJ and normalized to the size of the medulla (Figure 4U). VEGFR2 signal intensity was measured within equal square ROIs within cortex and medulla domains across different timepoints. Apoptosis and macrophage accumulation were measured as caspase-3^+^ or IBA1^+^ pixel area normalised to the total adrenal area (area%) after applying a threshold in Fiji/ImageJ to generate a binary mask. Volumetric view of phagocytic macrophages and apoptotic cells in Figure 6K-M was obtained with Arivis 4D software.

Statistical analyses were performed with GraphPad Prism V6.01 software. Unpaired t test was used to compare control vs. mutant conditions, (ns) p>0.05; (*) p<0.05; (**) p<0.01; (***) p<0. 001 (****) p<0.0001.

### Single-cell RNA-seq dataset analysis

The gene expression matrices provided by Hanemaaijer *et al*, 2021 and Furlan *et al*, 2017 were downloaded from GEO database (GSE162238, GSE99933) and analysed with the *Seurat R package* v4.0.1 (Butler *et al*, 2018). All functions mentioned in this paragraph are from the *Seurat R package* and are used with the default parameters unless otherwise noted. The dataset from Hanemaaijer *et al*. includes 12 samples −6 developmental time points (from E13.5 to P5), in duplicate. Furlan *et al*. dataset includes 2 samples corresponding to E12.5 and E13.5 stages. All samples were normalized with the *SCTransform function* from the *sctransform R packages* (Hafemeister & Satija, 2019), then combined using *harmony integration workflow* (Korsunsky *et al*, 2019). On Hanemaaijer *et al*. dataset, we reported an average of 2791 reads/cell and 1460 genes/cell, while Furlan *et al*. dataset shows an average of 269559 reads/cell and 6978 genes/cell. Principal component analysis was performed with *RunPCA* and the number of principal components that were retained (nPCs = 20) was selected after inspection with *ElbowPlot*. Cell clustering was performed with *FindNeighbors* and *FindClusters* at a resolution of 0.3. Clusters were characterized according to the expression of known chromaffin, cortex, endothelial, immune, stroma, SCP, bridge, neuroblast signature genes. Single-cell expression levels for Plexin-D1, Neuropilin1 and 2, Sema3C were plotted on UMAPs with the *FeaturePlot function*.

## Notes

### Competing Interest Statement

The authors have declared no competing interest.

### Summary of Updates

Corrections in the method section; author list updates.

