## Supplementary Figures for "A neurovascular template guides the spatial and functional compartmentalization of the adrenal gland"

### Figure S1

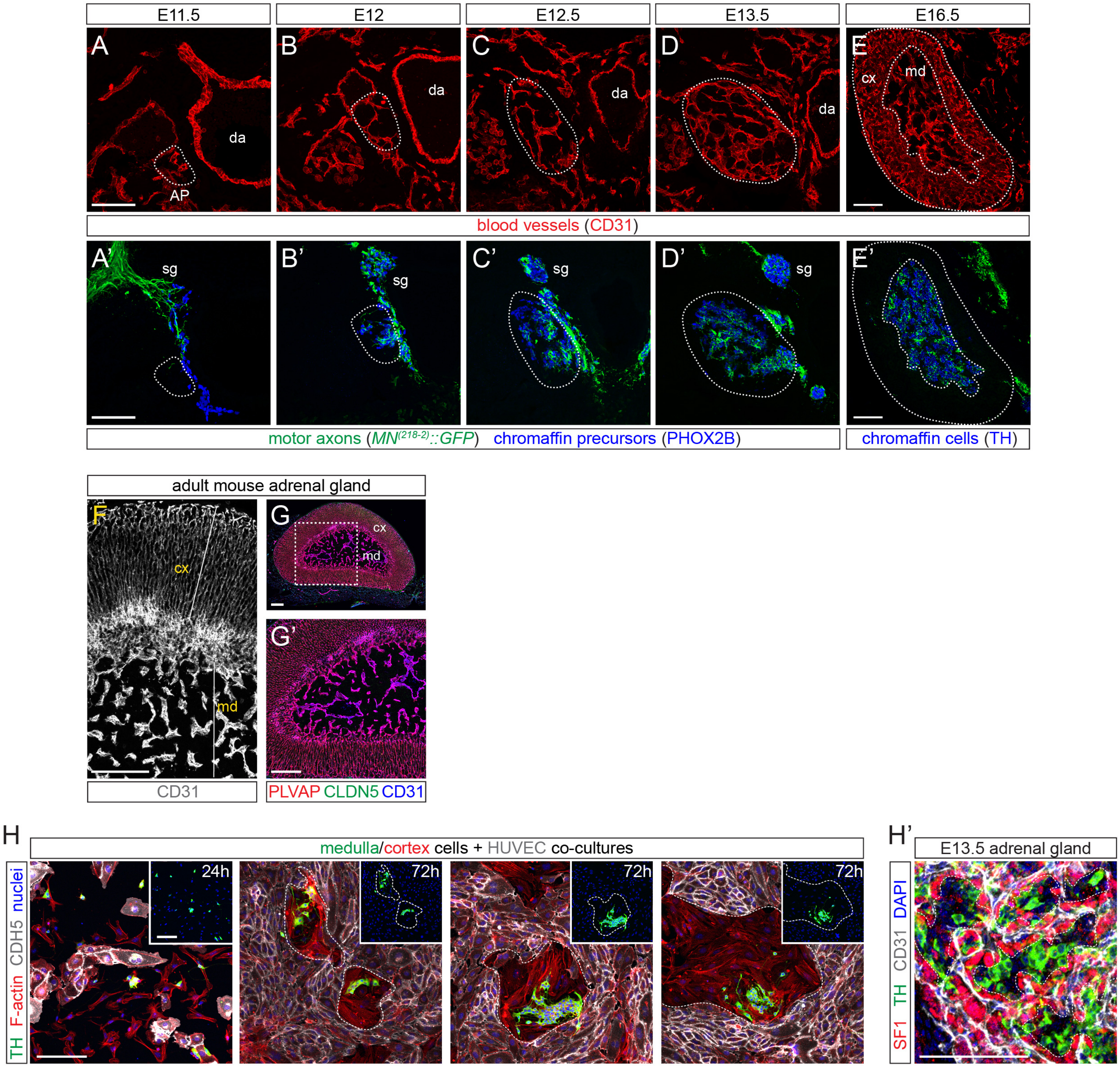

E11.5
E12
E12.5
E13.5
E16.5
A
B
C
D
E
da
AP
cx
md
blood vessels (CD31)
A'
B'
C'
D'
E'
sg
motor axons (MN(218-2)::GFP) chromaffin precursors (PHOX2B)
chromaffin cells (TH)
adult mouse adrenal gland
F
G
G'
CD31
PLVAP CLDN5 CD31
H
medulla/cortex cells + HUVEC co-cultures
24h
72h
TH F-actin CDH5 nuclei
H'
E13.5 adrenal gland
SF1 TH CD31 DAPI

**Suppl Figure 1: Neurovascular assembly during adrenal gland morphogenesis. (A – E')** Separate channels from adrenal gland developmental time-course shown in Figure 1G – K. Blood vessels are identified with anti-CD31 labelling (A – E). In A' – E', motor axons are labelled by *$MN^{(218\text{-}2)}$::GFP* (green), while chromaffin cells by either PHOX2B or TH (blue). **(F – G')** Adult adrenal gland sections immunostained with CD31 in grey (F) or with the vessel permeability marker PLVAP in red, the barrier marker Claudin5 (CLDN5) in green and CD31 in blue (G, G'). **(H, H')** Adrenal gland cell-HUVEC co-culture assay showing progressive redistribution of chromaffin ($TH^+$, green) and cortical cells (identified by Phalloidin staining, PHALL, red) dissociated from E14.5 adrenal glands of wild-type embryos, relative to HUVECs ($CDH5^+$, grey), at 24 and 72 hours after plating. DAPI identifies cell nuclei. H' shows *in vivo* distribution of cortical ($SF1^+$, red) and medullary cells ($TH^+$, green) relative to adrenal vessels ($CD31^+$, grey) in a wild-type E13.5 embryo. AP: adrenal primordium; cx: cortex; da: dorsal aorta; md: medulla; sg: sympathetic ganglion. Scale bars: A – E', 100 µm; F – G', 200 µm; H, H', 100 µm.

### Figure S2

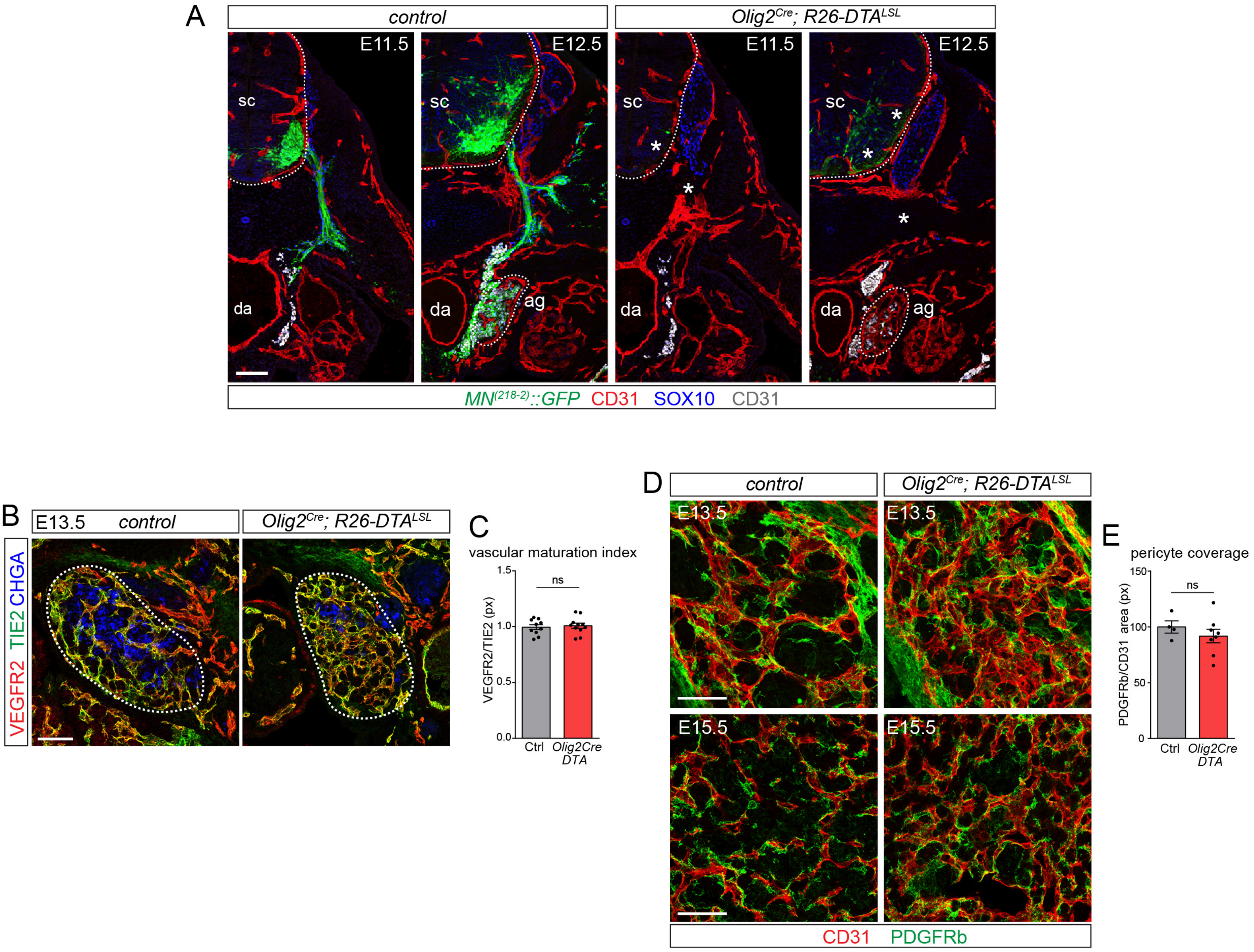

A
control
Olig2Cre; R26-DTALSL
E11.5
E12.5
sc
da
ag
MN(218-2)::GFP CD31 SOX10 CD31
B
E13.5 control
Olig2Cre; R26-DTALSL
VEGFR2 TIE2 CHGA
C
vascular maturation index
VEGFR2/TIE2 (px)
ns
Ctrl
Olig2Cre DTA
D
control
Olig2Cre; R26-DTALSL
E13.5
E15.5
CD31 PDGFRb
E
pericyte coverage
PDGFRb/CD31 area (px)
ns
Ctrl
Olig2Cre DTA

**Suppl Figure 2: Medullary signals influence adrenal vessel density but not maturation. (A)** Transverse embryo sections showing MN loss (asterisks) in E11.5 and E12.5 *Olig2*$^{Cre}$*; R26-DTA*$^{LSL}$ (right) compared to controls (left). MNs are marked by *MN*$^{(218\text{-}2)}$*::GFP* (green), CD31 labels endothelial cells (red), PHOX2B identifies sympathetic neurons and chromaffin cells (grey), SOX10 labels SCPs (blue). The spinal cord (sc) and adrenal gland (ag) are outlined. **(B)** Adrenal gland sections from E13.5 *Olig2*$^{Cre}$*; R26-DTA*$^{LSL}$ embryos (right) and controls (left). VEGFR2 (red) marks angiogenic vessels, while TIE2 (green) is expressed by the mature vasculature. Chromogranin A (CHGA) labels chromaffin cells. **(C)** Ratio between angiogenic (VEGFR2$^{+}$) and mature (TIE2$^{+}$) vessel area in the adrenal gland of *Olig2*$^{Cre}$*; R26-DTA*$^{LSL}$ versus controls at E12.5 and E13.5. **(D)** Adrenal sections from E13.5 (top) and E15.5 (bottom) *Olig2*$^{Cre}$*; R26-DTA*$^{LSL}$ embryos (right) and controls (left) immunostained for CD31 (red) and the pericyte marker PDGFRβ (green). **(E)** Pericyte coverage measured as the ratio between PDGFRβ and CD31 areas in E13.5 and E15.5 *Olig2*$^{Cre}$*; R26-DTA*$^{LSL}$ and control embryos. ag, adrenal gland; da, dorsal aorta; sc, spinal cord. Scale bars: Scale bar: A, 100 µm; B, 50 µm; D, 50 µm.

Figure S3

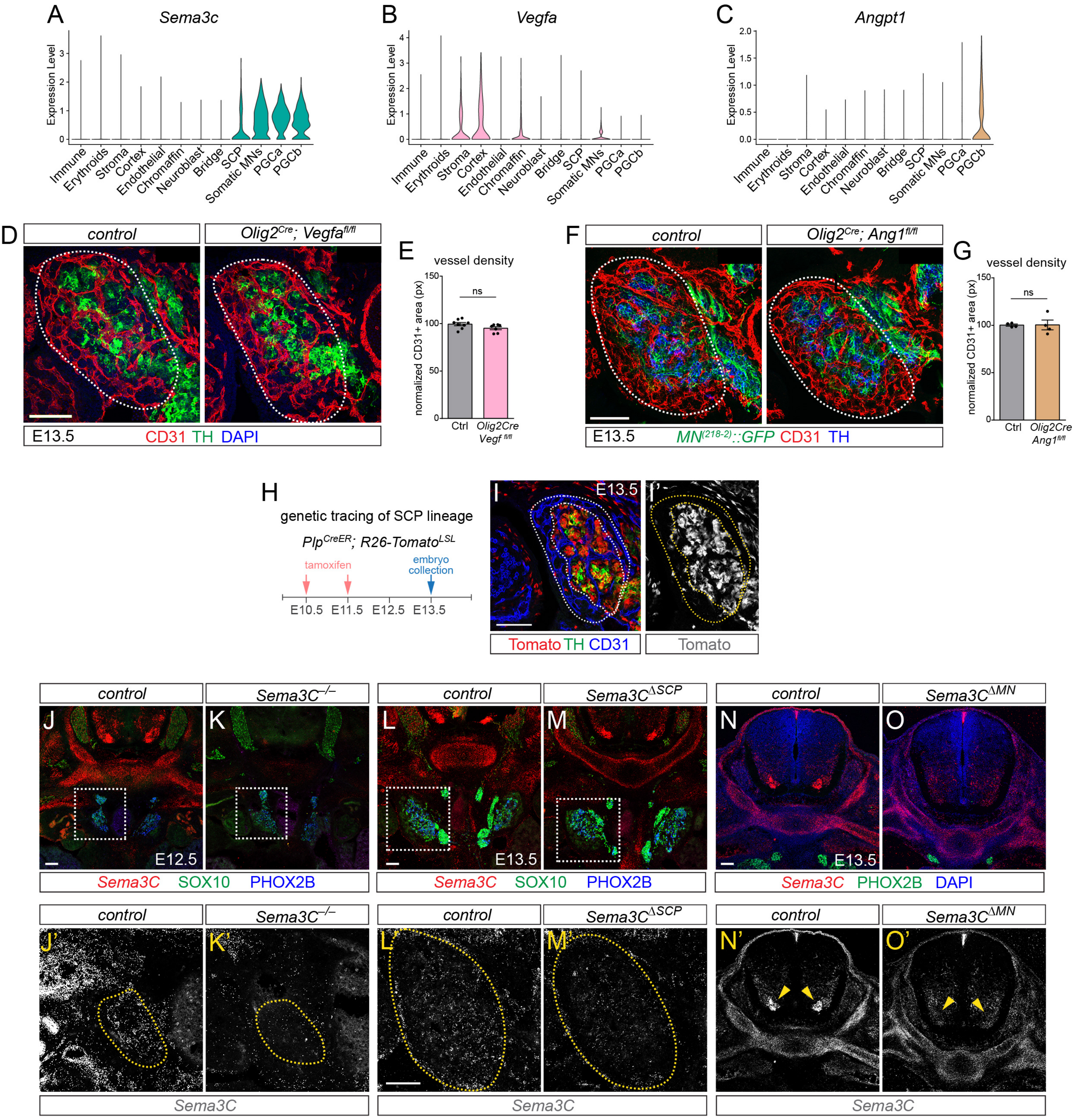

A
Sema3c
Expression Level
Immune
Erythroids
Stroma
Cortex
Endothelial
Chromaffin
Neuroblast
Bridge
SCP
Somatic MNs
PGCa
PGCb
B
Vegfa
C
Angpt1
D
control
Olig2Cre; Vegfafl/fl
E13.5
CD31 TH DAPI
E
vessel density
normalized CD31+ area (px)
ns
Ctrl
Olig2Cre Vegf fl/fl
F
control
Olig2Cre; Ang1fl/fl
E13.5
MN(218-2)::GFP CD31 TH
G
vessel density
normalized CD31+ area (px)
ns
Ctrl
Olig2Cre Ang1fl/fl
H
genetic tracing of SCP lineage
PlpCreER; R26-TomatoLSL
tamoxifen
embryo collection
E10.5
E11.5
E12.5
E13.5
I
I'
E13.5
Tomato TH CD31
Tomato
control
Sema3C–/–
control
Sema3CΔSCP
control
Sema3CΔMN
J
K
L
M
N
O
E12.5
E13.5
E13.5
Sema3C SOX10 PHOX2B
Sema3C SOX10 PHOX2B
Sema3C PHOX2B DAPI
J'
K'
L'
M'
N'
O'
Sema3C
Sema3C
Sema3C

**Suppl Figure 3: Loss of nerve-derived Sema3C, but not Vegfa or Angiopoietin-1, leads to vascular abnormalities. (A - C)** Violin plots showing the expression of *Sema3C* (A), *Vegfa* (B) and *Angpt1* (C) among adrenal gland cell types and spinal motor neuron populations from integrated datasets shown in Figure 3A and 3B. **(D)** Adrenal gland (outlined) from E13.5 *Olig2$^{Cre}$; Vegf $^{Fl/Fl}$* (right) and controls (left) immunostained for TH (green) and CD31 (red). DAPI labels nuclei (blue). **(E)** Adrenal vessel density in *Olig2$^{Cre}$; Vegf $^{Fl/Fl}$* and controls at E13.5. **(F)** Adrenal gland (outlined) from E13.5 *Olig2$^{Cre}$; Ang1$^{Fl/Fl}$* (right) and controls (left). Visceral motor axon terminals are labelled with *MN$^{(218-2)}$::GFP* (green), TH identifies chromaffin cells (blue) and CD31 labels vessels (red). **(G)** Adrenal vessel density in *Olig2$^{Cre}$; Ang1$^{Fl/Fl}$* and controls at E13.5. **(H)** Schematic of activation of the *Plp$^{CreER}$* for targeting of the SCP lineage. **(I, I')** Adrenal gland sections immunostained for TH (green) and CD31 (blue) in E13.5 *Plp$^{CreER}$; R26-TOM$^{LSL}$*. Tamoxifen injection at E10.5 and E11.5 activates Tomato expression (red) in nearly all medullary chromaffin cells. Tomato signal is shown separately in the right panels (grey). The adrenal cortex and medulla are outlined. **(L, K')** *Sema3C* (red) expression, detected by RNAscope, is significantly reduced in E12.5 *Sema3C$^{-/-}$* (K, K') but not in control embryos (J, J'). SOX10 (green) labels SCPs, PHOX2B (blue) identifies immature chromaffin cells. The boxed areas are magnified in J' and K' showing *Sema3C* in grey. The adrenal gland area is outlined. **(L, M')** RNAscope shows a reduction of *Sema3C* expression (red) selectively in the adrenal gland of *Sema3C $^{ΔSCP}$* (M, M') compared to controls (L, L'). SOX10 (green) labels SCPs, PHOX2B (blue) identifies immature chromaffin cells. L' and M' are magnified insets from the dotted squared area showing expression of *Sema3C* (grey) in the adrenal gland (outlined). **(N, O')** RNAscope shows selective MN ablation of *Sema3C* (red) in E13.5 *Sema3C $^{ΔMN}$* (O, O'), but not controls (N, N'). PHOX2B (green) identifies sympathetic ganglia, DAPI labels nuclei. *Sema3C* (grey) is shown separately in N' and O'. Yellow arrowheads point to spinal motor columns. Scale bars: 100 µm

### Figure S4

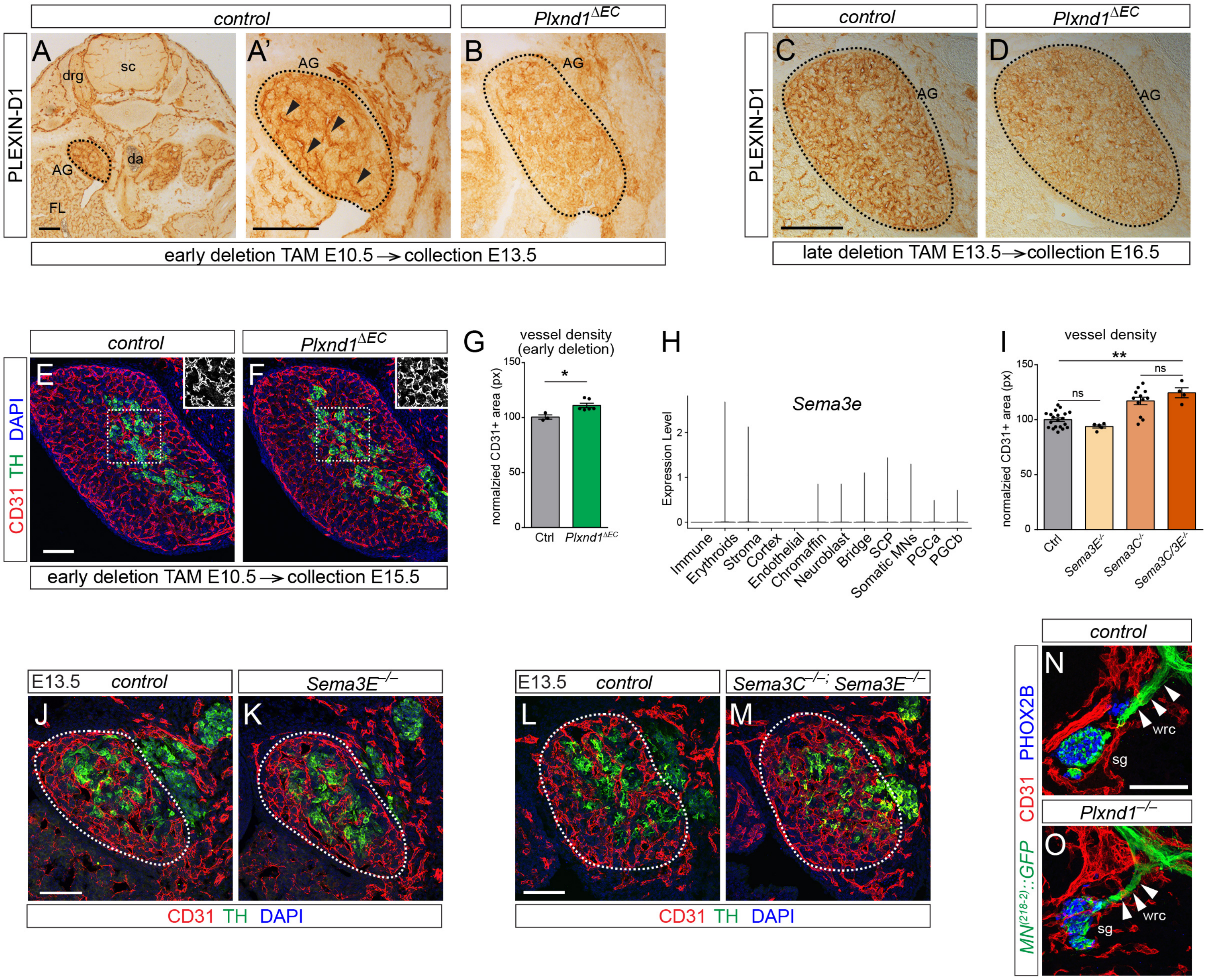

control
Plxnd1ΔEC
PLEXIN-D1
A
A'
B
drg
sc
AG
da
FL
early deletion TAM E10.5 → collection E13.5
control
Plxnd1ΔEC
C
D
AG
late deletion TAM E13.5 → collection E16.5
control
Plxnd1ΔEC
E
F
CD31 TH DAPI
early deletion TAM E10.5 → collection E15.5
G
vessel density (early deletion)
normalzied CD31+ area (px)
150
100
50
0
*
Ctrl
Plxnd1ΔEC
H
Sema3e
Expression Level
2
1
0
Immune
Erythroids
Stroma
Cortex
Endothelial
Chromaffin
Neuroblast
Bridge
SCP
Somatic MNs
PGCa
PGCb
I
vessel density
normalzied CD31+ area (px)
150
100
50
0
**
ns
ns
Ctrl
Sema3E-/-
Sema3C-/-
Sema3C/3E-/-
E13.5
control
Sema3E−/−
J
K
CD31 TH DAPI
E13.5
control
Sema3C−/−; Sema3E−/−
L
M
CD31 TH DAPI
control
Plxnd1−/−
N
O
wrc
sg
PHOX2B
CD31
MN(218-2)::GFP

**Suppl Figure 4: Adrenal vessels sense repulsive medullary Sema3C through endothelial Plexin-D1. (A – D)** Transverse embryo sections from E13.5 (A – B) and E16.5 (C, D) controls (A, A', C) and *Plxnd1* $^{\Delta EC}$ (B, D) immunostained for Plexin-D1. The adrenal gland is outlined. Three consecutive tamoxifen injections from E10.5 (early gene deletion, A – B) or E13.5 (late gene deletion, C, D) efficiently ablate Plexin-D1 from endothelial cells in *Plxnd1* $^{\Delta EC}$, but not in littermate controls. **(E, F)** Adrenal gland sections of E15.5 *Plxnd1* $^{\Delta EC}$ (F) and controls (E) upon early endothelial deletion of *Plxnd1* (tamoxifen administration from E10.5). TH identifies chromaffin cells (green), CD31 blood vessels (red), DAPI cell nuclei (blue). Insets at the top-right corner show medullary vessels ($CD31^+$) in grey. **(G)** Adrenal vessel density in E15.5 *Plxnd1* $^{\Delta EC}$ and controls upon early endothelial deletion of *Plxnd1*. **(H)** Violin plot showing *Sema3E* expression in adrenal gland cell types and spinal motor neuron populations from integrated datasets shown in Figure 3A and 3B. **(I)** Adrenal vessel density in control, *Sema3E* $^{-/-}$, *Sema3C* $^{-/-}$ and compound *Sema3C/3E* knock-outs at E13.5. Data points from *Sema3C* $^{-/-}$ and littermate controls are also shown in Figure 3P. **(J – M)** Adrenal gland sections of E13.5 *Sema3E* $^{-/-}$ (K) and compound *Sema3C* $^{-/-}$*; Sema3E* $^{-/-}$ mutants (M) compared to littermate controls (J, L) immunostained for TH (green) and CD31 (red). DAPI labels nuclei in blue. The adrenal gland is outlined. **(N – O)** Transverse E12.5 embryo sections showing white ramus communicans (wrc, arrowheads) (*MN* $^{(218\text{-}2)}$*::GFP* $^+$, green) innervating sympathetic ganglia ($PHOX2B^+$, blue) in *Plxnd1-/-* (O) and controls (N). CD31 is in red. AG, adrenal gland; da, dorsal aorta; drg: dorsal root ganglion; FL: fetal liver; sc, spinal cord; sg: sympathetic ganglion; wrc: white ramus communicans. Scale bars: A – D, 200 µm; E, F, 100 µm; J – M, 100 µm; N – O, 100 µm.

### Figure S5

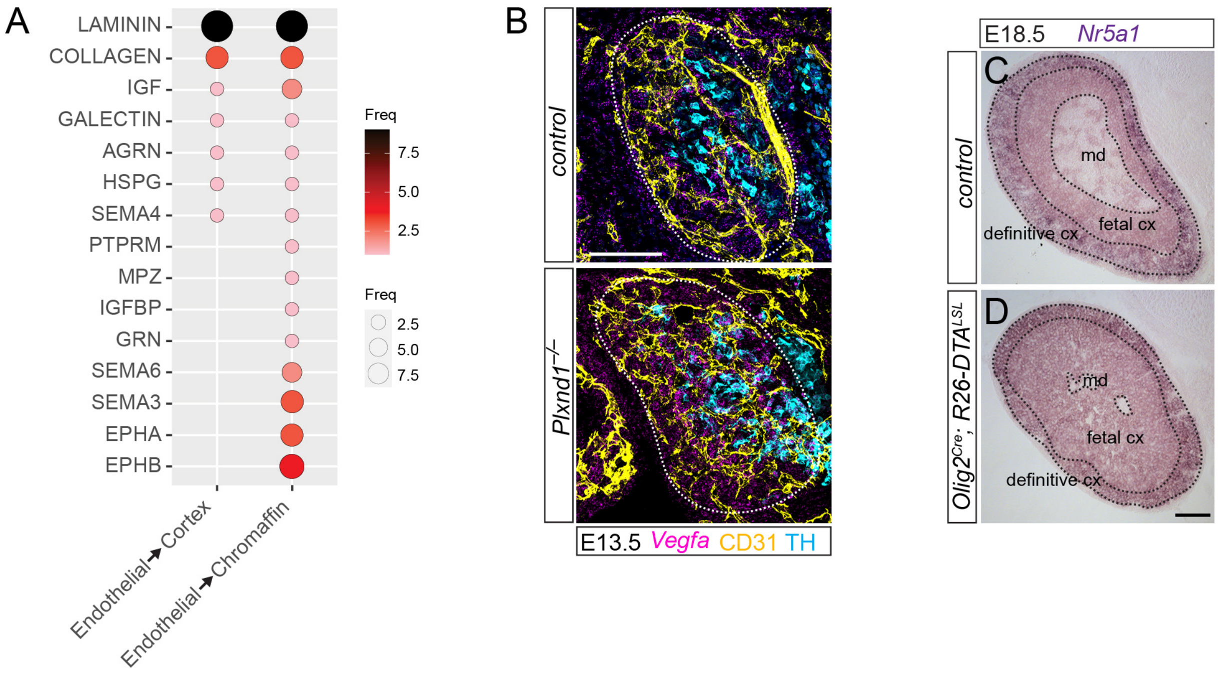
A
LAMINININ
COLLAGEN
IGF
GALECTIN
AGRN
HSPG
SEMA4
PTPRM
MPZ
IGFBP
GRN
SEMA6
SEMA3
EPHA
EPHB
Endothelial→Cortex
Endothelial→Chromaffin
Freq
7.5
5.0
2.5
Freq
2.5
5.0
7.5
B
control
Plxnd1−/−
E13.5 Vegfa CD31 TH
E18.5 Nr5a1
C
control
md
fetal cx
definitive cx
D
Olig2Cre; R26-DTALSL
md
fetal cx
definitive cx

**Suppl Figure 5: Adrenal endothelial cells recruit VEGF-expressing cortical cells. (A)** Dotplot showing the pathways mediating the Endothelial-to-Cortex and Endothelial-to-Chromaffin interactions in the developing adrenal gland using integrated datasets shown in Figure 3A. Dot size and color scale show the frequency of interactions. **(B)** *Vegf*a RNAscope (magenta) on E13.5 control (top) and *Plxnd1-/-* (bottom) adrenal glands (outlined) immunostained for TH (cyan) and CD31 (yellow). Magnified insets without TH staining are shown in Figure 5S, T. **(C, D)** *In situ* hybridization for the cortical marker *Nr5a1* on adrenal sections from E18.5 controls (C) and *Olig2*$^{Cre}$*; R26-DTA*$^{LSL}$ (D). The medulla, as well as the fetal and definitive cortex, are outlined. Scale bars: 100 µm.

### Figure S6

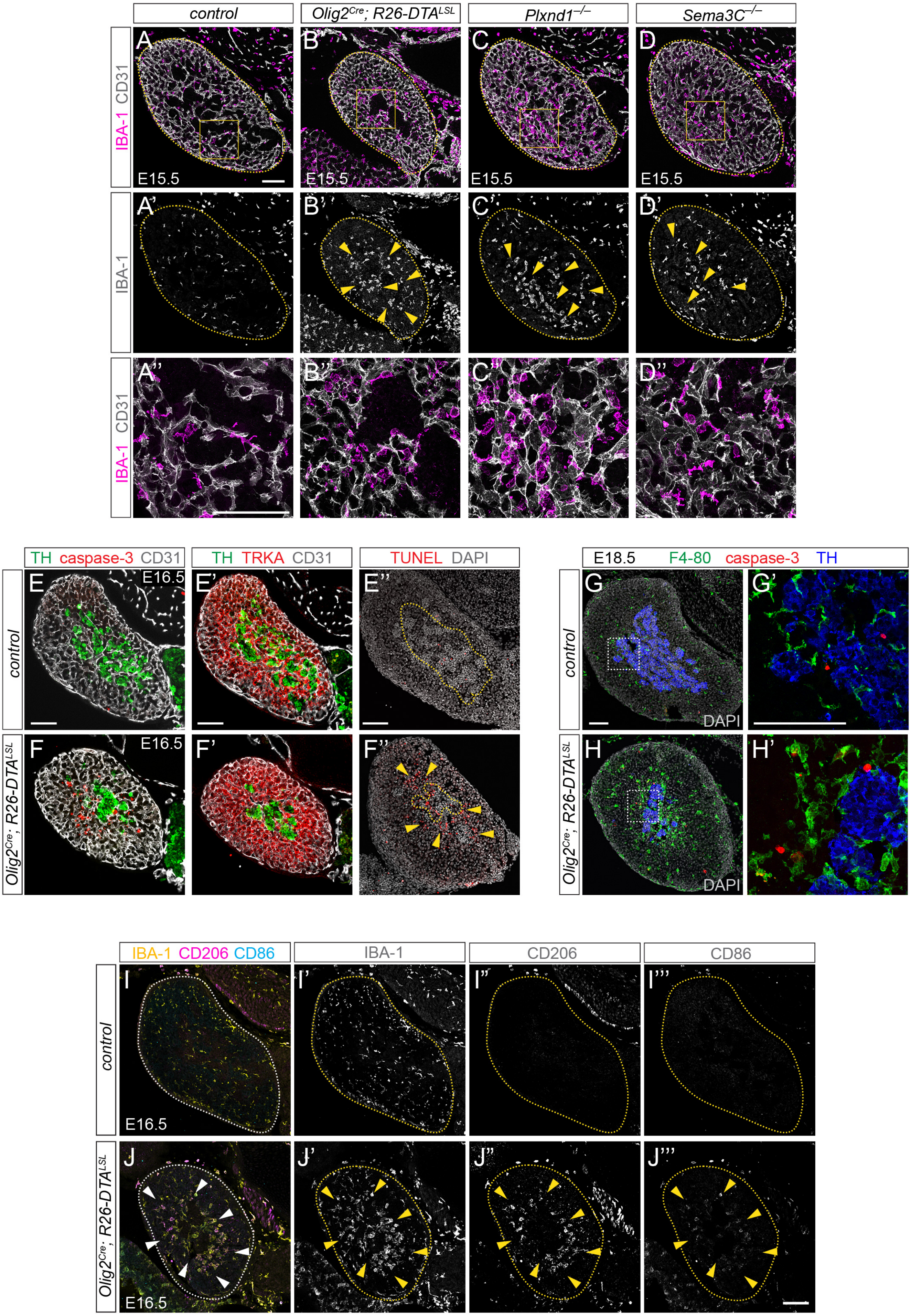

control
Olig2Cre; R26-DTALSL
Plxnd1−/−
Sema3C−/−
IBA-1 CD31
IBA-1
E15.5
TH caspase-3 CD31
TH TRKA CD31
TUNEL DAPI
E16.5
E18.5 F4-80 caspase-3 TH
DAPI
IBA-1 CD206 CD86
IBA-1
CD206
CD86

**Suppl Figure 6: Phagocytic macrophages accumulate at the cortico-medullary border in mutants. (A – D'')** Adrenal sections from control (A – A''), *Olig2*$^{Cre}$*; R26-DTA*$^{LSL}$ (B – B''), *Plxnd1-/-* (C – C'') and *Sema3C -/-* (D – D'') embryos at E15.5. IBA1 (magenta) labels macrophages, CD31 (grey) blood vessels. A' – D' show IBA1 staining in grey. The adrenal gland shape is outlined. A'' – D'' show magnified insets from the boxed area in A – D. **(E – F')** Adjacent adrenal sections from *Olig2*$^{Cre}$*; R26-DTA*$^{LSL}$ (F – F') and control (E – E') immunostained for TH (green), CD31 (grey) and in red either caspase-3 for apoptosis (E, F) or TRKA to label cortical cells (E', F'). **(E'', F'')** TUNEL staining (red) on E16.5 *Olig2*$^{Cre}$*; R26-DTA*$^{LSL}$ (F'') and control (E'') adrenal gland sections. DAPI (grey) is used to identify the morphology of adrenal compartments (outlined). Arrowheads point to apoptotic cells accumulated at the cortico-medullary border **(G – H')** E18.5 adrenal sections from *Olig2*$^{Cre}$*; R26-DTA*$^{LSL}$ (H, H') and controls (G, G') immunostained for TH (blue), caspase-3 (red) and the macrophage marker F4-80 (green). DAPI in grey marks cell nuclei. Insets (G', H') show a magnified view of the boxed area. **(I – J''')** Adrenal sections from E16.5 *Olig2*$^{Cre}$*; R26-DTA*$^{LSL}$ (J) and controls (I) immunostained for the pan-macrophage marker IBA1 (yellow), the anti-inflammatory marker CD206 (magenta) and the pro-inflammatory marker CD86 (cyan). Merged view as well as single separate channels are shown. The adrenal gland shape is outlined. Arrowheads show anti-inflammatory macrophage accumulation in mutants. Scale bars: 100 µm.
